# A Nanomule Peptide-siRNA Conjugate that Traverses the Intact Blood Brain Barrier and Attenuates Stroke

**DOI:** 10.1101/871186

**Authors:** Brett A. Eyford, Chaahat S.B. Singh, Thomas Abraham, Lonna Munro, Kyung Bok Choi, Rhonda Hildebrandt, Tracy Hill, Ian Welch, Mark Okon, Timothy Z. Vitalis, Reinhard Gabathuler, Jacob A. Gordon, Hans Adomat, Emma S.T. Guns, Chieh-Ju Lu, Cheryl G. Pfeifer, Mei Mei Tian, Wilfred A. Jefferies

**Affiliations:** Michael Smith Laboratories, University of British Columbia, 2185 East Mall, Vancouver, BC, Canada, V6T 1Z4. 13; The Vancouver Prostate Centre, Vancouver General Hospital, 2660 Oak Street, Vancouver, BC, Canada, V6H 3Z6. 16; Centre for Blood Research, University of British Columbia, 2350 Health Sciences Mall, Vancouver, BC, Canada, V6T 1Z4. 19; The Djavad Mowafaghian Centre for Brain Health, University of British Columbia, 2215 Wesbrook Mall, Vancouver, BC, Canada, V6T 1Z4. 22; Department of Medical Genetics, University of British Columbia, 2350 Health Sciences Mall, Vancouver, BC, Canada, V6T 1Z4. 25; Department of Neural and Behavioral Sciences and Microscopy Imaging Core Lab, Pennsylvania State College of Medicine, 500 University Drive, Hershey, PA, USA, 17033-2390. 28; Centre for Comparative Medicine, University of British Columbia, 4145 Wesbrook Mall, Vancouver, BC, Canada, V6T 1W5 31; Department of Chemistry, University of British Columbia, 2350 Health Sciences Mall, Vancouver, BC, Canada, V6T 1Z4. 34; Bioasis Technologies Inc., 14 Water Street, Guilford, CT 06437, USA.; Department of Urologic Sciences, University of British Columbia, Gordon & Leslie Diamond Health Care Centre, Level 6, 2775 Laurel Street, Vancouver, BC Canada V5Z 1M9 39; Department of Microbiology and Immunology, University of British Columbia, 2350 Health Sciences Mall, Vancouver, BC, Canada, V6T 1Z4. 42; Department of Zoology, University of British Columbia, 6270 University Blvd., Vancouver, BC, Canada, V6T 1Z4. 45

**Author notes:** Correspondence to Senior Author: Professor Wilfred A. Jefferies, Michael Smith Laboratories, University of British Columbia, 301-2185 East Mall, Vancouver, British Columbia, V6T 1Z4 Canada, and The Vancouver Prostate Centre, Vancouver General Hospital, 2660 Oak Street, Vancouver, BC, Canada, V6H 3Z6. These authors contributed equally to this work and are co-first authors (Brett A. Eyford, Chaahat S.B. Singh, Thomas Abraham).

## Abstract

The blood-brain barrier (BBB), hinders the distribution of therapeutics, intended for treatment of diseases of the brain. A twelve-amino acid peptide, termed MTfp, was derived from MTf, and retains the ability to cross the BBB intact and ferry cargo into intacellular organelles within neurons, glia and microglia in the brain. A novel MTfp-siRNA peptide-oligonucleotide conjugate (POC), directed against NOX4, a gene known to potentiate ischemic stroke, was chemically synthesized. The MTfp-NOX4 siRNA POC traversed the BBB, resulting in the knockdown of NOX4 expression in the brain. Following induction of ischemic stroke, animals treated with the POC exhibited significantly smaller infarcts; accompanied by significant protection against neurological deterioration and improved recovery. The data demonstrates that the MTfp portion, of this novel POC, can facilitate BBB transcytosis; where the siRNA moiety can elicit effective therapeutic knockdown of a gene associated with a disease of the central nervous system (CNS). This is a general platform to transport therapeutics to the CNS and thereby, offers new avenues for potential treatments of neuropathologies that are currently refractory to existing therapies.

## Introduction

Many neurological diseases of the central nervous system (CNS) are completely untreatable because the Blood-Brain Barrier (BBB) excludes efficacious drugs from entering the brain^1–5^. Methods developed to enhance the delivery of drugs to treat diseases in the brain often fail to provide significant improvements to long-term survival^6–16^. Widely hailed methods of delivery to the brain often successfully demonstrate transport of the carrier or localization in an area of the brain after the blood-brain barrier is disrupted by pathogenesis or design but few, if any actually treat a CNS disease when the blood-brain barrier remains impermeable. Previous approaches include delivery of micro-encapsulated drugs^3–5^ and radical methods to transiently increase the permeability of the BBB, allowing diffusion of injected drugs from the periphery into the brain^17, 18^ The latter approach has the added toxicity caused by uncontrolled entry of the blood constituents into the brain and vice-versa. Traditionally, drug design has had to consider factors such as lipid solubility, charge, molecular weight and antiport action^19^. Drugs conjugated to small hydrophobic peptides and proteins^4, 20^ or antibodies^21^, able to bind to receptors expressed on the luminal surface of the BBB, have been studied. For example, MRC OX26, an antibody made against the transferrin receptor (TfR)^22^ has been used to deliver drugs to tumours in the periphery^22^ and in the brain^23^. Although delivery to the brain is sometimes achievable, its use is limited due to saturation of the receptor, low dissociation rate of the antibody on the abluminal side of the BBB and consequential recycling of the receptor back to the blood^24^. Hyper-immunity against the carrier may also limit repeated treatments with the same drug conjugates. In addition, many of the targeted receptors are widely expressed in other tissues resulting in potential toxicity^22^. Thus, novel approaches are required to increase the survival of patients with CNS tumours and other currently untreatable brain diseases such as lysosomal storage diseases.

It has been found that antibodies conjugated to drugs can cross the BBB as a result of their interaction with specific receptors, which suggests that drugs conjugated to the natural, endogenous ligands for these receptors may be of value in the delivery of systemic-borne therapeutic agents to the brain^5^. In order to address this, we have investigated the expression and distribution of ligands and receptor molecules on brain capillary endothelium^25, 26^ and these may provide novel routes of entry into the brain. We have previously shown that that melanotransferrin (MTf; Uniprot P08582. Also known as p97, MFI2 and CD228), a mammalian iron-transport protein, is an effective carrier for delivery of drug conjugates across the BBB into the brain^25–29^. MTf is actively transported across the BBB, by receptor mediated transcytosis at rates 10 to 15 times higher than those obtained with either serum transferrin (Tf) or lactoferrin (Lf)^25^. In addition, therapeutic drugs conjugated to MTf, can be shuttled across the BBB. For example, mice suffering from otherwise inoperable brain tumours were treated with MTf-drug conjugates resulting in reduction of tumour growth^30^. MTf is established as an endogenous protein with clear potential as a BBB drug delivery vehicle. However, MTf is a relatively large (738 residue, ∼80 kDa), iron-binding glycoprotein that can be expressed in both soluble and glycosylphosphatidylinositol-anchored forms^31^. The variability in conformation, glycosylation, anchoring and metal binding, presents potential complications to the use of MTf as a robust, reproducible, clinically useful drug delivery vehicle. Therefore, to improve upon the utility of MTf as a drug-delivery vector, we fragmented the protein in order to identify the minimally active region that retains the capability of carrying molecular cargo across the BBB. Here we report the identification of a fully functional, twelve amino acid, BBB carrier peptide, termed MTfp.

Unfortunately, most candidate BBB transcytotic carriers fail in the ultimate test of therapeutic efficacy in the CNS. Therefore, in a proof-of-concept study, we have focused on measuring treatment in a model of ischemic stroke as a test of this new, transcytotic peptide vector. Stroke is one of the leading causes of death in North America. It is caused by the impairment of cerebral blood flow (ischemic stroke) or the rupture of blood vessels in the brain (haemorrhagic stroke). The interruption of oxygenation and metabolites results in neuronal death. The utility of short antisense/interfering RNA (siRNA) therapeutic approaches for CNS diseases such as stroke has been limited by their biodistribution *in vivo*. On their own, siRNA preferentially localize to the kidney and liver, and their exclusion from the brain continues to hamper their potential and this represents a significant technical hurdle^32^. However, if brain delivery was possible, we hypothesized that reducing the expression of genes in the brain that potentiate the pathophysiology of stroke could be neuroprotective.

NADPH oxidase type 4 (NOX4; Uniprot Q9JHI8) primarily produced by brain-resident microglia was chosen because this protein is known to be up-regulated during acute ischemic stroke, it is a major source of oxidative stress leading to neuronal apoptosis. Animals deficient in NOX4 are strongly protected from ischemic stroke^33, 34^ and its expression lies within the brain; beyond the BBB. Therefore treatments that alter NOX4 expression must penetrate the intact BBB, thereby validating them as being delivered by transcytosis. To this end, a novel peptide-oligonucleotide conjugate^35^ (POC) composed of MTfp and NOX4-targeted siRNA was *de novo* synthesised and assessed for improving outcomes in a model of ischemic stroke. Data support the utility of MTfp targeted delivery to the CNS as a useful therapy to ameliorate the neuropathology associated with ischemic stroke and potentially other diseases affecting the brain.

## Results

### Identification of MTfp

In order to identify a fragment of MTf, also termed, p97, MFI2 and CD228 (or commercially as *Transcend*) that retains the ability to cross the BBB, purified, recombinant human MTf was digested with either hydroxylamine (NH_2_OH), cyanogen bromide (CNBr) or trypsin. The resulting fragments were placed on the luminal side of an *in vitro* bovine BBB model and, after two hours, the presence of MTf fragments on the abluminal side was assessed by SDS-PAGE (NH_2_OH and CNBr) and mass spectrometry (all digestions). No fragments from the CNBr or NH_2_OH digests were found on the abluminal side of the BBB model (data not shown). However, 50 potential tryptic peptides were tested (Supplemental Table 1), and five were identified in the abluminal medium of the *in vitro* BBB model: DSSHAFTLDELR; ADTDGGLIFR; VPAHAVVVR; ADVTEWR; and YYDYSGAFR. To determine relative transport efficiencies, these five peptides were synthesized and retested individually in the same BBB model system. Quantitation of transcytosis was achieved by spiking known concentrations of stable isotope labeled peptides prior to mass spectrometry. The most efficiently transcytosed peptide (DSSHAFTLDELR, MTf_460-471,_ now known as MTfp, or commercially as xB^3^ ^TM^ peptide) was selected as the most promising candidate for use as a BBB transport vector (Table 1). MTfp transit across the BBB *in vivo* was tested in mice by injecting a MTfp-Cy5 conjugate intravenously (IV), followed by visualization and quantification by 3D deconvolution fluorescence microscopy (Fig. 1 and Supplemental Table 2). Approximately two times greater fractional fluorescence was measured in the brain parenchyma of MTfp-Cy5 injected wild type (WT) mice when compared to control mice injected with PBS or reversed MTfp-Cy5 (revMTfp-Cy5). Interestingly, Tfp penetrated the BBB with efficiency similar to that of a larger, known brain-targeting peptide derived from a rabies virus glycoprotein (RVGp) a virus known to enter the brain^36, 37^. The MTfp was shown to colocalize in neurons, microglia and astrocytes in the CNS. As shown in Figure 2, demonstrated that 90% of the neurons expressing NeuN co-labelled with MTfp; 70% of astrocytes expressing GFAP co-labelled with MTfp; and 80% of microglia expressing TMEM co-labelled with MTfp. The subcellular markers investigated show that 40% of the MTFp is found in the lysosomes and approximately 20% is found in the early endosomes. This data demonstrates for the first time that MTFp injected IV, is widely dispersed in the brain and it is taken up into intracellular organelles.

**Figure 1.**
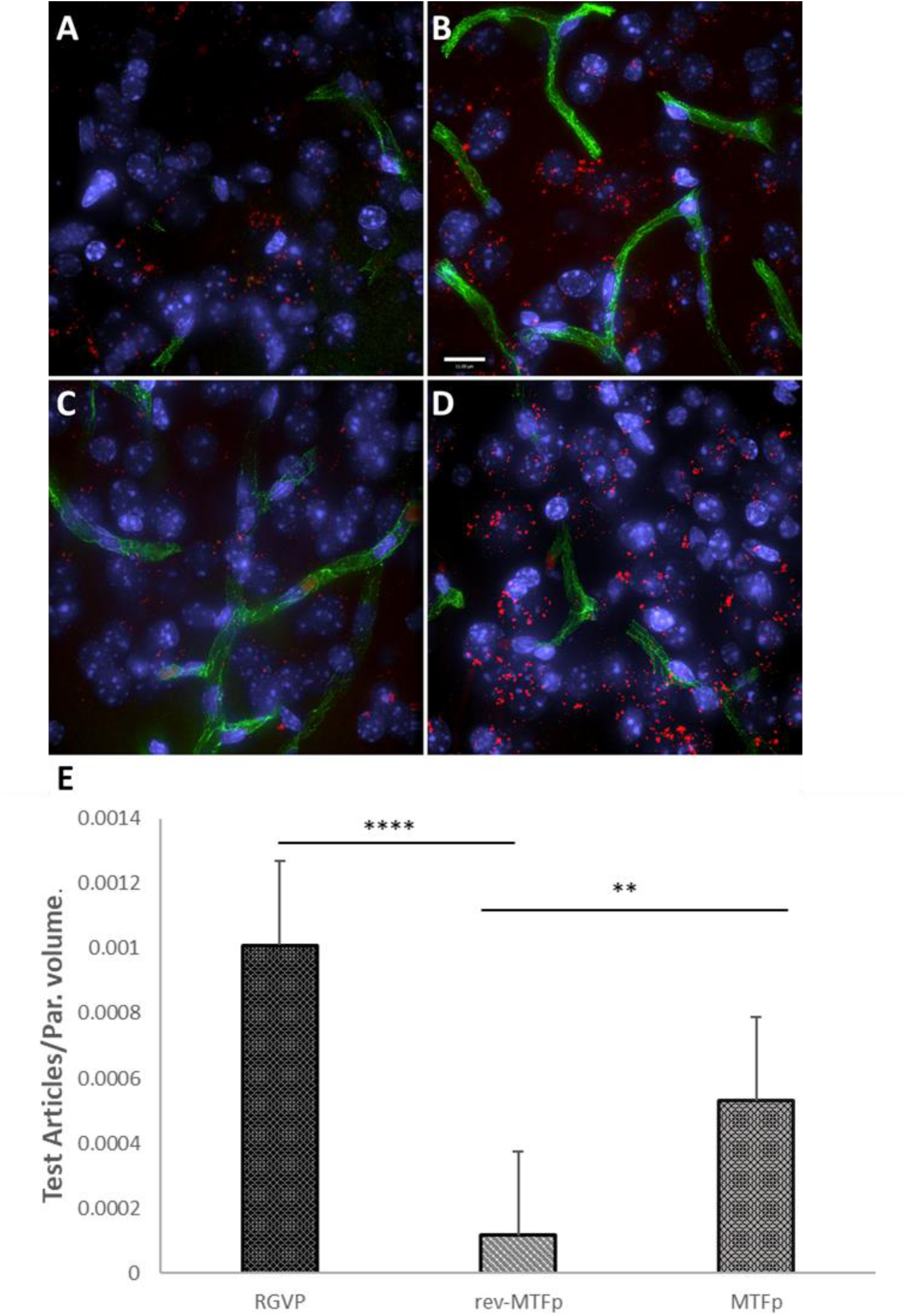
Representative deconvolved images showing localization of MTfp in the brains of mice. Cell nuclei are blue (DAPI) and capillaries are green (FITC). [**A**] Fluorescence (red) in the brain for a mouse treated with PBS (*i.e.* background fluorescence); [**B**] Cy5 fluorescence in the brain after IV injection with RVGp-Cy5; [**C**] Cy5 fluorescence in the brain after IV injection of revMTfp-Cy5; [**D**] Cy5 fluorescence in the brain after IV injection of MTfp-Cy5; [**E**] Brain distribution of MTfp in wild type mice. Values indicate total volume of Cy5 fluorescence in each tissue normalized to tissue volume (VTA_BPV_ in Supplemental Table 2) and normalized by subtracting the background fluorescence seen in mice treated with PBS. Data are represented as means ± SD (n=3, 3 sections/animal and 4-11 fields of view per section).

**Figure 2:**
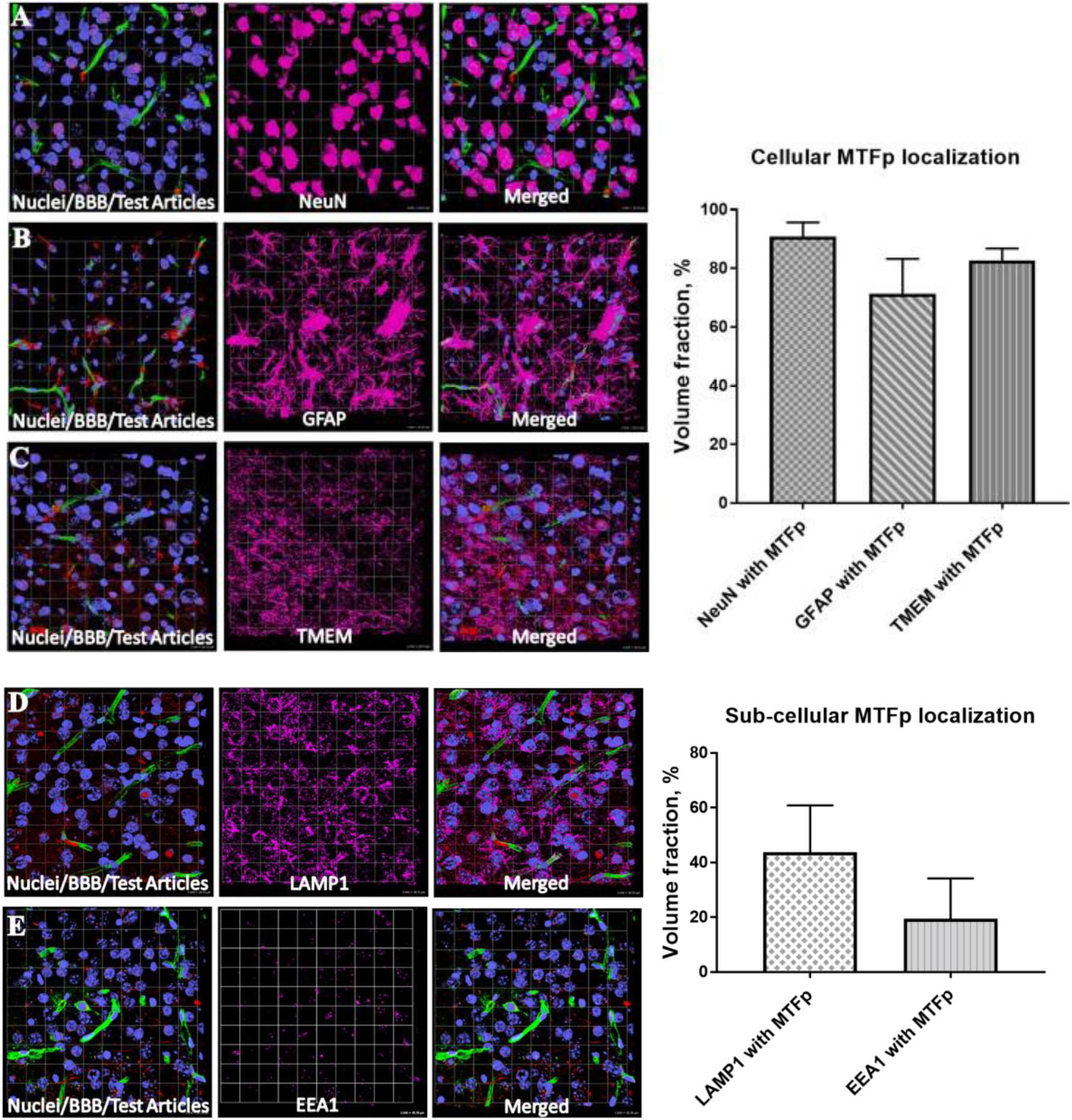
Representative 3D confocal images showing localization of various cell types and subcellular objects with MTfp in the mouse brain sections. Cell nuclei are in blue color (DAPI) and BBBs are in green color (FITC). **A, B and C; cellular localization of MTFp and D, E and F; subcellular localization of MTFp.** [**A**] Localization of neuronal marker NeuN (Pink) with MTfp (Red); [**B**] Localization of Astrocytes marker GFAP (Pink) with MTfp (Red); [**C**] Localization of Microglia marker TMEM (Pink) with MTfp (Red); [**D**] Localization of Lysosomal marker LAMP1 (Pink) with MTfp (Red); [**E**] Localization of Endosomal marker EEA1 (Pink) with MTfp (Red). The graphs represent the extent of localization of various cell types and subcellular objects with MTfp in the CNS as a percentage of fluorescence volume fraction.

**Table 1.**
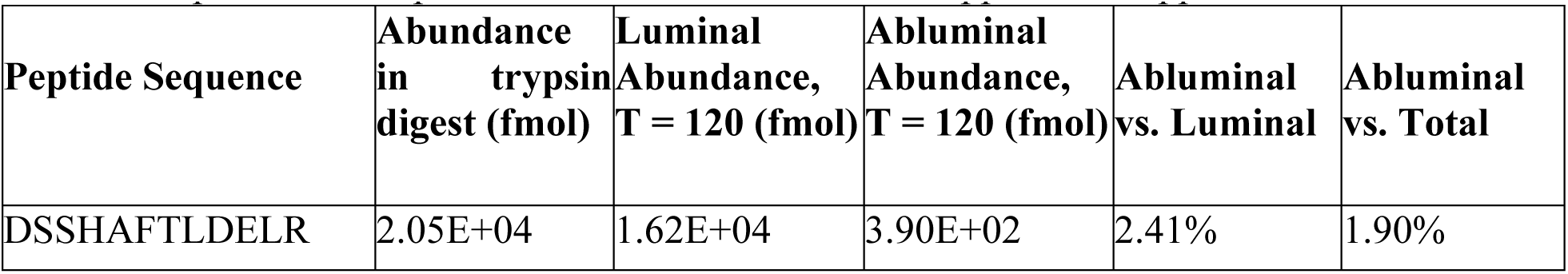
A MTf Tryptic peptide crosses the BBB in vitro. A model of the BBB was created to screen for MTf peptides that cross the BBB *in vitro*. Molar abundances are normalized against compartment volumes is used as the measure of transcytosis efficiency. The discrepancy between initial concentration and total recovered concentration is likely explained by a combination of cellular uptake, peptide degradation and adhesion to surfaces and matrices. A comprehensive summary of the screen for the fraction of peptides found in the abluminal compartment compared to the initial concentration appears in Supplemental Table 1.

### Tissue distribution: Recovery of the intact MTfp in the brain and other tissues

We developed a methodology to quantitate the intact MTFp peptide within the brain and other tissues. Initially, MTFp peptide proved quite difficult to extract from tissue matrices. Very poor recoveries resulted with simple clarified homogenates using ultrafiltration, SPE, solvent precipitation or combinations thereof. Urea solubilizing on its own resulted in slight improvement. Inclusion of considerable acetic acid ultimately enhanced extraction efficiency enough for a usable assay, albeit still suboptimal. Use of stable isotope labeled standards SIS is essential with R^2^<0.99 for calibration in brain homogenate with good alignment with non-matrix prepared samples. The assay is feasible with analytical flow triple quad (QQQ) but substantially superior with the nanoflow Orbitrap (OT) platform in terms of both selectivity and sensitivity but with less robust retention time alignment. The triply charged 3+ ion was more intense and fragmented more efficiently with the OT and provided a better endpoint as opposed to the 2+ with the QQQ, however, further optimizing with the OT might improve the 2+ response. Detection limits are approximately 20ng/g for triple quad and sub ng/g for Orbitrap with this assay system. Run times in the order of the analytical flow are likely possible with somewhat larger bore nLC tubing and columns packed in house with little loss in sensitivity and further improvements extraction may be possible as well, both of which will be followed up in future work. The levels of MTfp in the brain, liver and kidney following IV dosing (100 µl of a 1mM solution tail vein), as determined for the 30min time point is shown in Figure 3 based on the protocol described by Bickel^38^. At the 30-minute time point the brain concentration obtainedwas 64 nM. An approximating of the initial rate of transport into the brain is therefore 2 nM/min. At the 30-minute time point, the liver and kidney concentration obtained was 220 nM and 300 nM respectively, translating to an initial rate of transport of 7.3 nM/min for liver and 9.9 nM/min for kidney. Associated representative chromatograms for MTfp are summarized in Figure 3B. These mass-spectrometry based studies conclusively demonstrate that MTFp is transported intake into these tissues.

**Figure 3.**
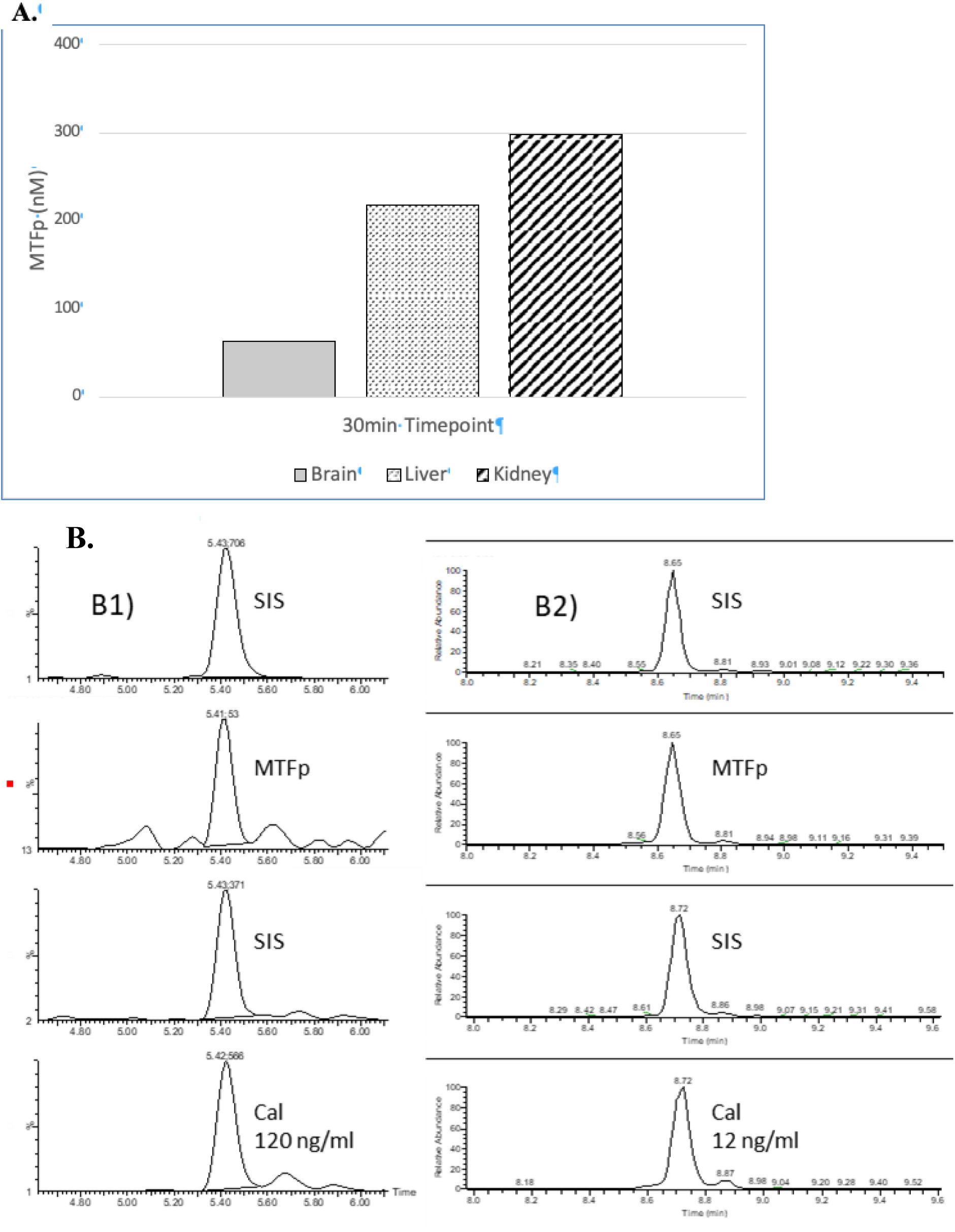
Biodistribution of MTfp. [**A**] Biodistribution; concentration of MTfp and its 837 distribution in different organs at 30-minute time point in CD1 mice treated with 4.7mg/kg. For 838 brain; n=3, mean ± SEM; single measurements with other tissues. 2.1 nM/minute was seen as the 839 initial rate of MTFp entry into the brain, calculated as the slope of the time vs concentration. **[B]** 840 Selected chromatograms highlighting LCMS tissue analysis. **B1)** brain from 30-minute treated 841 animal and 120ng/ml equivalent spiked blank brain homogenates. 10µl sample inject volume with 842 0.3ml/min flow, 2.1×100mm column, QQQ. **B2)** Same sample along with 12ng/ml calibration, 843 2µl sample inject volume with 300nl/min flow, 50µm x 15cm PepMap RSLC C18, OT.

### MTfp-NOX4 siRNA reduces stroke damage

To assess whether MTfp could deliver RNA cargo to the CNS, fluorescent MTfp was conjugated to NOX4-specific siRNA and injected IV into mice. The brains were then examined by 3D confocal fluorescence microscopy (Fig. 4 and Supplemental Table 3). The conjugation of MTfp to siRNA allowed increased localization to the brain parenchyma compared to siRNA alone (Fig. 4). NOX4 knockdown was measured by real-time polymerase chain reaction (RT-PCR). MTfp-siRNA facilitated reduction in NOX4 mRNA to 20% of the levels observed with the PBS injection control (Fig. 5A). Interestingly, injection of NOX4-specific siRNA alone resulted in reduction of expression to 45% of control levels. We believe that this may be due to the siRNA acting upon the vascular endothelium but not the brain parenchyma, since siRNA’s do not cross the BBB.

**Figure 4.**
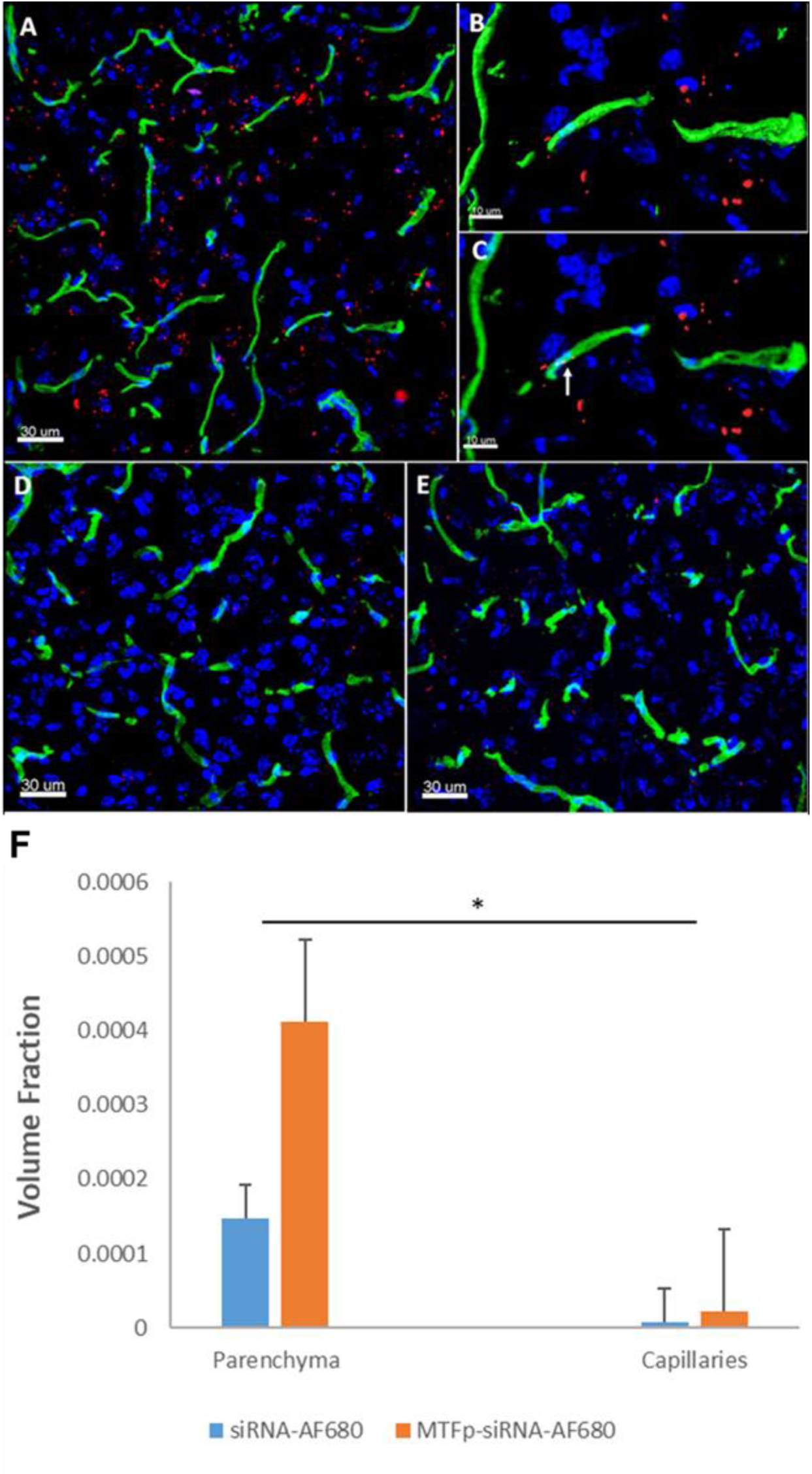
Representative 3D confocal images showing localization of MTfp-siRNA in the brain of wild type mice. Cell nuclei are blue (DAPI) and capillaries are green (Tomato lectin-FITC). **[A]** AF680 fluorescence (red) in the brain of a mouse treated with siRNA-MTfp_AF680_; **[B]** Shows the surface rendered (quantified) FITC labelled capillaries (green) and siRNA-MTfp_AF680_ (red); **[C]** Example of siRNA-MTfp_AF680_ localized with the blood capillaries (yellow, see arrow); **[D]** AF680 fluorescence (red) in the mouse brain treated with PBS (*i.e.* background fluorescence); **[E]** AF680 fluorescence (red) in the mouse brain treated with siRNA-AF680. **[F]** Brain distribution of MTfp-siRNA in wild type mice. Values indicate total AF680 fluorescence normalized to total 859 tissue volume (VTA in Supplemental Table 3) and then normalized to the total AF680 fluorescence 860 seen in PBS (background). Data are represented as means ± SD (n=3, 8 fields of view per animal). 861 *P-value < 0.05.

**Figure 5.**
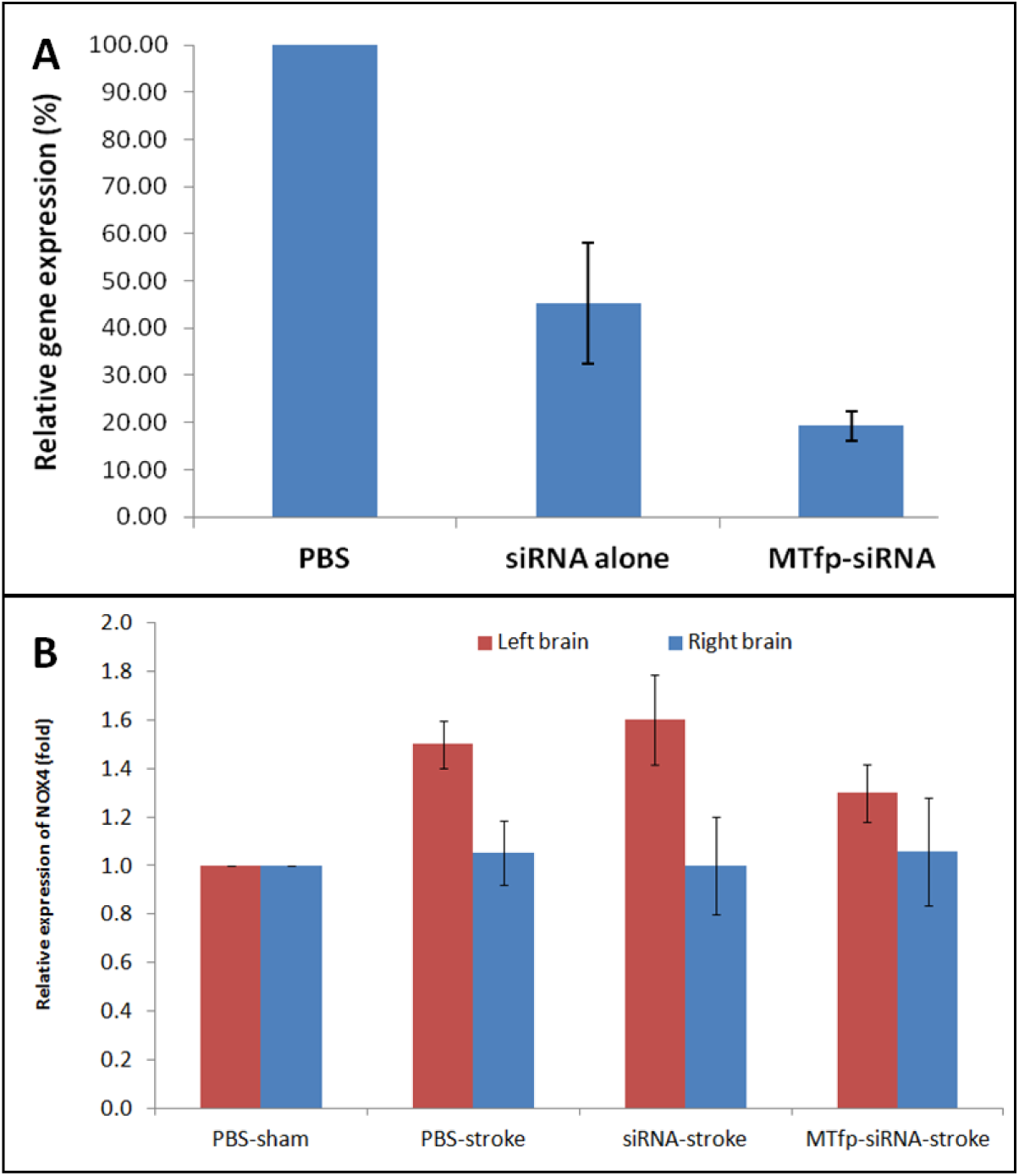
NOX4 mRNA expression in the brain after siRNA treatment and stroke induction. **[A]** Relative gene expression of *Nox4* in the whole brain of WT mice was quantified by RT-PCR (GAPDH as a reference gene). **[B]** Relative gene expression of *Nox4* in the mouse brain hemispheres by RT-PCR following treatment and 24 hours post-stroke (GAPDH as a reference gene). For each treatment group, the *Nox4* expression in the left hemisphere (stroke) is normalized against the right hemisphere (contralateral) of the same animal. Data are shown as mean ± SEM.

To demonstrate the therapeutic efficacy of cargo delivery to the CNS by MTfp, MTfp conjugated to NOX4-specific siRNA was examined in mouse model of ischemic stroke. Mice were dosed three times, at one-hour intervals, and then subjected to ischemic stroke in the left hemisphere for a duration of one hour by middle cerebral artery occlusion (MCAO). We chose to administer treatment (MTfp-siRNA, MTfp-scramRNA, NOX4 siRNA alone and PBS controls) prior to stroke induction since the integrity of the BBB could be compromised by the invasive nature of stroke induction. Furthermore, this therapy is intended to reduce NOX4/hypoxia associated pathology and will have no effect if blood flow to the site of injury is impaired (*i.e.* blood clots or filament occlusion). Animals were sacrificed at 24 hours post reperfusion. It is known that induction of ischemic stroke leads to elevated CNS expression of NOX4^33^. This trend is seen in our experiments as well when stroke is induced in animals that were treated with PBS, NOX4 siRNA alone (Fig. 5B) or MTfp-scramRNA control (Supplemental Fig. 1 and 2). However, when treated with MTfp-siRNA, NOX4 up-regulation was significantly prevented (Fig. 5B). Finally, we assessed whether MTfp-siRNA mediated NOX4 knockdown resulted in protection against neurological deterioration. Mice were treated and subjected to ischemic stroke as described above. Neuroscore^39^ was measured 0.5h after removal of anesthesia (to determine surgical efficiency) and again at 24 hours post stroke (to measure recovery and outcome). After the second behavioural assessment, animals were sacrificed and infarct volume was quantitated by 2,3,5-triphenyltetrazolium chloride (TTC) staining. Total infarct volume as well as motor deficits were measured (Fig. 6 and Supplemental Table 4). Compared to PBS controls, sham surgery siRNA alone and MTfp-scramRNA controls, the MTfp-siRNA resulted in significantly smaller infarcts (Fig. 6 and Supplemental Fig. 3) and corresponded to less severe paralysis immediately after cessation of MCAO (Fig. 6). Twenty-four hours after surgery, mice that had been treated with MTfp-siRNA achieved neurological scores of <1 which is functionally near normal, while the control animals remained mentally impaired.

**Figure 6.**
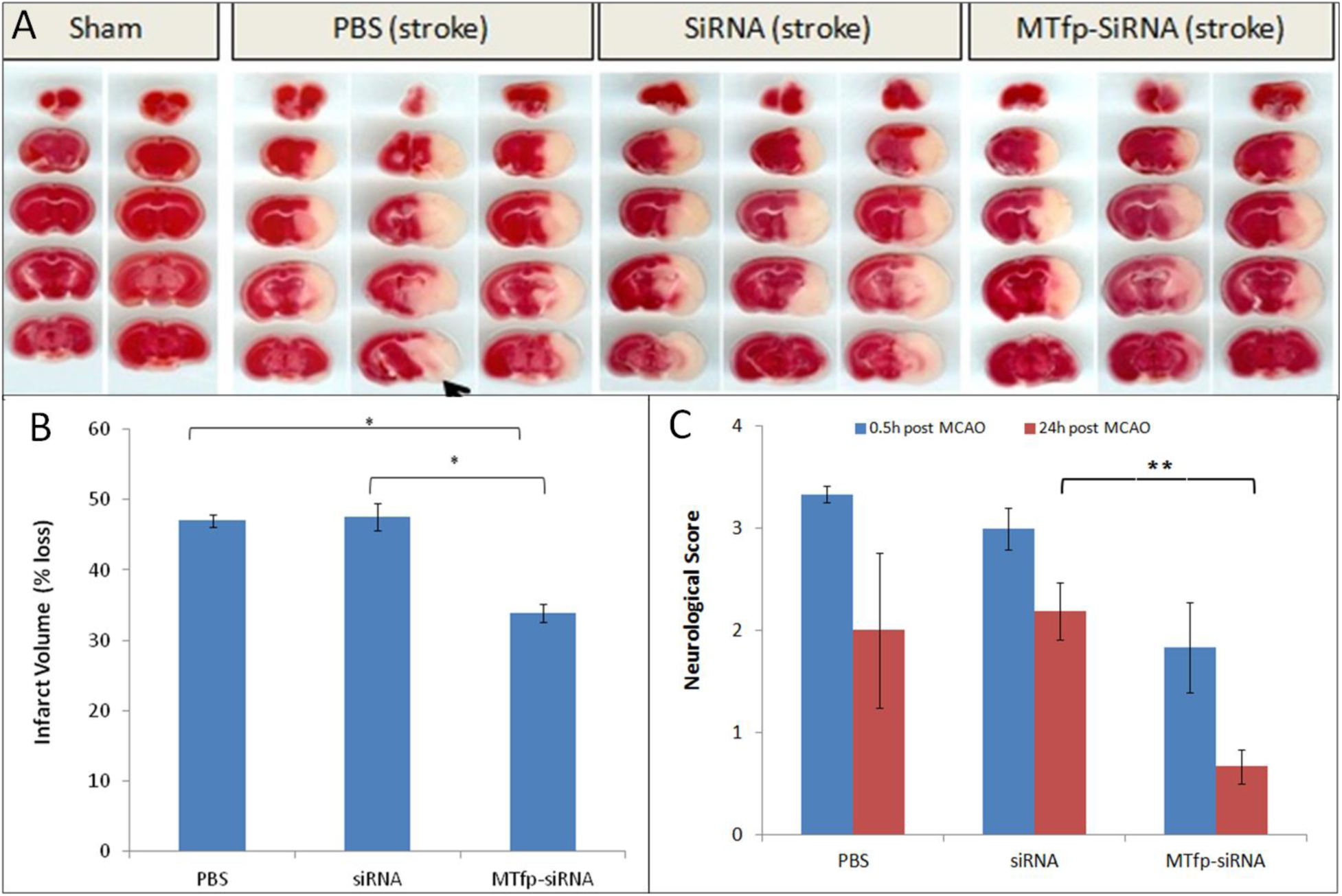
Treatment with MTfp-siRNA confers neuroprotection and reduces damage after ischemic stroke. **[A]** TTC-stained brain sections of mice receiving various IV injections, followed by ischemic stroke. Tissue was collected 24 hr after surgery **[B]** Infarct volume was quantitated by measuring absorbance of solvent extracted dye (*, *P<*0.05, one-way ANOVA). **[C]** Neuroscore at 0.5 hr and 24 hr after stroke induction in mice pretreated with siRNA, MTfp-siRNA or PBS control. \**P<*0.05, \*\**P<*0.001.

## Discussion

Designing efficient ‘vectors’ (antibodies, protein carriers, viruses, nanoparticles) to navigate and deliver therapeutics across the BBB, in a controlled and non-invasive manner, remains one of the key goals of drug development for brain diseases. Drug design for CNS diseases is constrained by factors such as lipid solubility, charge, molecular weight, and the antiport action of specific transporters^19^. Many methods developed to enhance the delivery of drugs to treat brain diseases have failed to provide significant improvements to long-term survival^6–16^. Drugs conjugated to small hydrophobic peptides, proteins^20^, or antibodies^21^ are able to bind to receptors expressed on the luminal surface of the BBB, the accumulation in the brain resulting efficacious effects on disease is severely lacking. Unfortunately, these approaches are often limited due to saturation of the receptors, low dissociation rate on the abluminal side of the BBB, and recycling of the receptor back to the lumen^24^. In addition, many targeted receptors are widely expressed in tissues other than the brain, resulting in potential toxicity^22^ and reduced transport to the brain because of competition in peripheral tissues. We have previously shown that mammalian iron-binding protein melanotransferrin is able to deliver small anti-cancer agents to the brain and reduce the growth of tumours therein resulting in extending the survival of the treated animals. However, there are limitations in using MTf and other large protein carriers. These include; heterogeneity in post-translational modifications, variablity in the site of cargo chemical conjugation and the adoption of different structural conformations upon conjugation that may alter the targeting moiety or the action of the drug. The other major problem is that approaches using chemical conjugates are not scalable and batch production requires standards of equivalency that are cost and time prohibitive. Here we report the identification of a twelve amino acid peptide (MTfp; DSSHAFTLDELR), derived from melanotransferrin protein, that retains the ability crosses the BBB *in vitro* and *in vivo*. Structural characterization of MTfp, by nuclear magnetic resonance, shows that the peptide is largely unstructured but possesses the propensity to adopt an alpha helical shape. This may facilitate binding to a candidate MTf receptor (Supplemental Discussion 1). We demonstrate that MTFp, injected IV, becomes widely dispersed within the brain where 90% of the neurons co-stain with MTfp; 70% of astrocytes co-stain with MTfp; and 80% of microglia co-stain with MTfp. Furthermore, we can track the MTFp into cells in the brain and find that 40% of the MTFp is found in the lysosomes and approximately 20% is found in the early endosomes. In order to assess if MTFp can be transported into the brain, we developed a new mass spectrometry methodology to quantitate the recovered MTFp within the brain. Using this novel LCMS technique, we successfully demonstrated that the undigested peptide is indeed present in tissues and this allowed us to calculate an initial transport rate for MTFp into the brain as 2nM/minute.

MTfp was synthesized as a POC with a specific siRNA intended to treat ischemic stroke, a common disease of the CNS. The resulting POC retains the peptide’s ability to access brain tissue where the siRNA is able to knockdown its target, resulting in efficacious outcomes in models of ischemic stroke. The relative distribution of MTfp-siRNA between the brain and the kidney and liver, where siRNAs generally localise, was not measured, however, it is clear that this novel POC directs its payload, at therapeutic levels, to the brain. This is a significant advantage over existing siRNA delivery systems.

Previous attempts to deliver siRNA to the CNS have relied on antibody-targeted liposomes^40^ or non-covalent, electrostatic interactions with a peptide derived from rabies virus glycoprotein (RVGp; used here as a positive control)^36, 37, 41^. The POC we describe is connected covalently, can be produced used standard peptide/RNA synthesis and conjugation technologies and is derived from a protein of human origin, reducing the likelihood of immunogenicity compared to a viral carrier.

We chose to focus on stroke for this proof-of-concept study because it is a leading cause of death worldwide, is difficult disease to treat and has a pathogenic gene as a contributing factor. Currently, recombinant tissue plasminogen activator is the only approved drug for treatment of stroke but due to contraindications, it can only be used in about 10% of patients. Thus, new and effective treatments are needed for stroke. Oxidative stress, which is the damage caused by reactive oxygen species, appears to play a central role in neuronal death in stroke. NOX4 is one of the major sources of oxidative stress and therefore an excellent therapeutic target in acute stroke. Upon ischemia, NOX4 is highly induced in both human and mouse brains. In addition, following cerebral ischemia, mice genetically deficient in NOX4 (Nox4^−/−^) are largely protected from oxidative stress, neuronal death and BBB leakage^33^. Furthermore, NOX4 is a culprit in CNS disease besides stroke and therefore this POC may have utility in other brain pathologies^42^.

Our data, obtained using a murine stroke model with the POC targeted against NOX4 for the amelioration of stroke appears to be the first example of efficacious/therapeutic gene knockdown in any model of a disease resident in the brain. In this proof-of-concept study we opted to deliver the treatment before stroke induction. This was done to show that MTfp can cross an intact BBB, without potential complications arising from disruption of the BBB during the stroke.

Alternatively, stroke impairs blood flow and it is possible that if therapies were to be administered IV during or after a stroke, they would be less likely to gain access to the affected tissues. We suspect that, to achieve best outcomes, the POC conjugate described here will have to be used in combination with traditional fibrolytic therapies. Despite the clinical limitations, the POC used in this study demonstrates that MTfp has the capability of facilitating access to the CNS, to a molecule (RNA) that would normally be excluded.

The use of a peptide as a BBB drug delivery platform, for a variety of CNS diseases, is intriguing. The peptide may be synthesised entirely *de novo,* granting greater control over issues associated with the stoichiometry of carrier to therapeutic and other inherent unpredictable problems of production such microheterogeneity due to unwanted by-products of conjugation to larger proteins. Furthermore, the nucleotide sequence coding for MTfp could be incorporated into recombinant protein therapies, intended for treatment of CNS disease (e.g. monoclonal antibodies or enzyme replacement therapies).

In conclusion, this work identifies a novel, non-invasive methods of introducing effective therapeutic compounds across the intact BBB by transcytosis and into neurons, glia and microglia within the brain thereby allowing the potential treatment of many neuropathologies. Future focus will be on expanding the utility of this new carrier to traverse therapeutics across the battlement of the BBB for the treatment of other pathologies within the physiological fortress of the CNS.

## Methods

### Animal experiments

All protocols and procedures involving the care and use of animals in these studies reviewed and approved by the Ottawa-National Research Council of Canada (NRC) or University of British Columbia, Animal Care Committees.

### Purification and fragmentation of MTf

Recombinant human MTf was expressed and purified at the University of British Columbia, as described previously^29, 43^. To perform trypsin digestion, 200 µL of MTf (10 mg/mL) were diluted in 1 mL of 0.15 M Tris, 6 M guanidine, 10 mM dithiothreitol, pH 9. The sample was heated at 60°C for 15 minutes and cooled to room temperature. The solution was further diluted with 1.8 mL of 0.15 M Tris, pH 9. Forty µg of sequencing grade modified trypsin (Cat. #V5111, Promega) were dissolved in 400 µL of 0.15 M Tris and added to the reaction mixture (1:50 trypsin to MTf ratio). The sample was incubated at 37°C for 20h, lyophilized and stored at −20°C. The digest was dissolved in 5 mL of 0.1% formic acid and the tryptic peptide mixture was cleaned using Sep-Pack C8 cartridges (Cat. # WAT036775, Waters). The tryptic peptides were lyophilized and stored at −20°C until transcytosis assays were performed.

### *In vitro* BBB model and transcytosis assays

Work involving peptide transcytosis through the *in vitro* BBB model was performed by Cellial Technologies (Lens France). The *in vitro* model of the BBB consists of primary bovine brain capillary endothelial cells grown to confluence on collagen coated; polycarbonate transwell inserts (0.4 µm pore size, 24 mm diameter, Cat. # 3412, Corning) and supported by primary rat glial cells. Under these conditions, endothelial cells retain endothelial markers (factor VIII –related antigen, non-thrombogenic surface, production of prostacyclin, angiotensin converting enzyme activity) and characteristics of the BBB (presence of tight junctions and P-glycoprotein, paucity of pinocytotic vesicles, monoamine oxidase activity, γ-glutamyl transpeptidase activity and 500-800 Ω/cm^2^ electrical resistance)^44–46^.

Primary cultures of glial cells were isolated from newborn rat cerebral cortex^47^. After the meninges had been cleaned off, the brain tissue was forced gently through a nylon sieve. DMEM supplemented with 10% (v/v) fetal bovine serum, 2 mM glutamine and 50 µg/mL of gentamicin was used for the dissociation of cerebral tissue and development of glial cells. The glial cells were plated at a concentration of 1.25 x 10^5^ cells/mL in six-well plates and incubated at 37°C in 5% CO_2_. The medium was changed twice a week. Three weeks after seeding, cultures of glial cells become stabilized.

Endothelial cells were cultured on gelatin-coated Petri dishes in DMEM supplemented with 10% (v/v) calf serum, 10% (v/v) horse serum, 2 mM glutamine and 50 µg/mL gentamicin. One ng/mL of basic fibroblast growth factor was then added every other day. Under these conditions, endothelial cells form a confluent monolayer in 12 days. Transwell inserts were coated, on the upper side, with rat-tail collagen prepared according to the method of Bornstein^48^. Confluent endothelial cells were trypsinized and plated on the upper side of the filters at a density of 4 x 10^5^ cells per insert.

Lucifer Yellow (LY), at 20 µM, is used as a paracellular marker allowing the evaluation of the integrity of the BBB. Ringer-HEPES buffer (150 mM NaCl; 5.2 mM KCl; 2.2 mM CaCl_2_; 0.2 mM MgCl_2_•6H_2_0; 6 mM NaHCO_3_; 5 mM HEPES; 2.8 mM glucose) was added to the lower compartment (abluminal side) of a six-well plate (3 mL per well). One, mL Ringer-HEPES buffer containing the trypsin digest of MTf or synthetic peptide (0.25 mg/mL), in co-incubation with LY, was placed in the upper compartment (luminal side). Incubations were performed on a rocking platform at 37°C for 2h. Experiments were performed in triplicate with filters containing a confluent monolayer of endothelial cells or in triplicate with empty filters coated only with collagen. At the end of the incubation period, aliquots of abluminal and luminal liquid were collected. Detection of LY fluorescence was performed using a fluorescence counter (Fluoroskan Ascent, Thermolab Systems) to ensure BBB integrity. Supernatants from the luminal and abluminal compartments were lyophilized and stored at −80°C until peptide analysis by mass spectrometry.

### Mass Spectrometry screen

Peptide identification was performed by MRM Proteomics (Victoria, British Columbia). Lyophilized supernatants from the *in vitro* BBB experiments were rehydrated with 2% acetonitrile, 0.1% formic acid. The samples were resolved by liquid chromatography at 0.4 mL/min, over 43 min, using an Agilent Eclipse Plus C18 (150 x 2.1 mm, 1.8 µm) column. The eluates were analyzed on an Agilent 6490 QQQ mass spectrometer in positive mode, controlled by Agilent’s MassHunter Workstation software (version B.04.01). All acquisition methods used the following parameters: 3500 V capillary voltage, 250 °C sheath gas temperature at 11 L/min sheath gas flow. The electron multiplier had a 400 V offset, the nebulizer was set to 30 p.s.i., and the first and third quadrupoles were set to unit resolution. A default fragmentor voltage and cell accelerator voltage of 380 V and 5 V, respectively. The maximum cycle time was 352.5 ms. All data was processed using Agilent Quantitation software (v.B.04.01) with automatic peak detection and smoothing Gaussian width of 5. All integrated peaks were manually inspected to ensure correct peak detection and integration.

For accuracy, peaks with intensity less than 50 counts and/or intra-ion pair retention time difference greater than 0.02 min were not quantitated. Quantitation was achieved by spiking a predetermined, optimized mixture of stable isotope standard peptides into the relevant experiment:

YYDYSGAFR = 500 fmol/µL; DSSHAFTLDELR = 2 pmol/µL; ADVTEWR = 100 fmol/µL; VPAHAVVVR = 200 fmol/µL; ADTDGGLIFR = 200 fmol/µL.

### Measurement of MTFp (DSSHAFTLDELR) distribution in tissues

In order to determine serum levels and associated brain levels of peptide following dosing, 6 mice were injected via the tail vein with 100 µl of a 1mM solution of peptide in PBS vehicle. At 30-minute time point (n=3) the animals were anesthetized by gradual induction to 5% isoflurane at a flow rate of 1 L/min oxygen and maintained via nose cone throughout the procedure. Brains were then removed from the mice and the isolated serum samples plus associated brain tissues were stored at −80°C prior to analysis by LCMS at the Vancouver Prostate Centre.

### MTFp (DSSHAFTLDELR) analysis by LCMS

Tissue samples were prepared by addition of water and an aliquot of 50X complete Protease inhibitor (Roche) to frozen brain samples (3X w/v) and homogenized using a Precellys tissue homogenizer (Bertin Technologies) with 3 x 45 second cycles at 6000 rpm. 50 mg or 90 mg aliquots were dispensed into Eppendorfs. Additional 40 µl water was added to the 50 mg samples followed by 2 µl of 1µg/ml MTfp SIS peptide (equivalent stable isotope labeled peptide, SISCAPA) and vortexing. Urea (60 mg) and acetic acid (40 µl) were then added with vortexing after each. Samples were placed in a sonication bath (VWR 150T) and sonicated approximately 30 minutes after which they were centrifuged for 5 minutes at 20,000 g. Supernatant was added to 10K Amicon filter units (0.5 ml) and centrifuged for 2 hours at 12,000 g. Flow through was taken through solid phase extraction (Waters C18 Vac Sep Pak 50 mg). Columns were equilibrated, each with 1 ml acetonitrile (ACN), (70% ACN/0.5% acetic) and 0.1% TFA (trifluoroacetic acid) and the entire 10K flow though was loaded. A 1ml 0.1% TFA and 0.1ml 0.5% acetic acid wash preceded elution with 0.5 ml 70% ACN/0.5% acetic acid. Eluate was concentrated down to 40-50 µl (Centrivap) prior to analysis. Calibration samples were prepared in an equivalent fashion using untreated brain homogenate, SIS and additionally including 2 µl MTfp (New England Peptide) with varying concentration.

Prepared samples were analysed at the Vanouver Prostate Centre, with a Waters Acquity/Quattro Premier as well as with an Easy n-LC 1200/Lumos by multiple reaction and parallel reaction monitoring (MRM, PRM). The Acquity was set up with a 2.1×100mm BEH C18, 1.7µm column at 35°C and a 0.2-9 minute 5-40% ACN gradient with 0.1% FA present with ramp to 95%, 2-minute flush and 2.5 minute re-equilibration for a 15 minute total run time. The Quattro was set with 3kV capillary, 120/300⁰C source/desolvation T, 900/50 L/hr N_2_ desolvation/cone gas, 8e-3 Ar collision gas, Q1 slightly below unit resolution. Ions selected were m/z 695.8 with 417.2, 893.4, 964.5 fragments (MTFp) and 700.8 with 427.2, 903.4, 974.5 fragments (SIS) with 35V cone and 35/32/30V collision respectively with 0.1 sec dwell each for both MRM sets. Inject volume was 10 µl and retention time was 5.45 minutes. Waters Quanlynx was used for raw data processing and exported to Excel for further processing.

The Easy nLC was set up with a PepMap RSLC C18 50 µm x 15 cm, 2µm column at 50⁰C and a 0-10 minute 4-30% ACN gradient followed by 12minute 80% ACN flush. Column equilibration was 4 µl and sample injects 2 µl with 6 µl loading volume with total run times about 40 minutes and MTFp eluting at about 8.5 minutes. The Easynano source was operated at 1900V and m/z 375-1500 orbitrap (OT) scan data collected at 60,000 resolutions, RF lense 30%, AGC 4e5, 50 ms max. MS^2^ data was collected with quad isolation m/z 0.8, HCD activation 30% (target ions m/z 695.8335, 464.5606 for MTFp; 700.8643, 467.9144 for SIS), OT 30,000 resolution m/z150-1500, RF 60%, AGC 5e4, 54 sec max. MS^2^ OT data was processed with Thermo Quan Processor/Browser software selecting m/z 964.45, 532.25, 974.5, 542.3 as respectively as fragments for PRM with 0.2Da mass windows and exported to Excel as with the MRM data. The initial rate of entry was calculated as the slope of the time vs concentration graph plotted with the 0 and 30-minute time points. The time-point and concentrations observed in the brain were used from the LCMS analysis. 2.1 nM/minute was the initial rate of MTFp entry into the brain and the initial rates were determined by the protocol described by Bickel^38^.

### Synthesis of MTf and MTfp conjugates

MTfp (DSSHAFTLDELRYC), reversed MTfp (revMTfp; RLEDLTFAHSSDYC), and RVGp (YTIWMPENPRPGTPCDIFTNSRGKRASNGYC) were synthesized by Anaspec (Fremont USA) and conjugated to Cy5. Each peptide was synthesized with addition ‘YC’ residues on the C terminus to facilitate fluorescent labelling. After the reaction, the crude reaction mixture was purified using semi-preparative reverse phase C18 chromatography. The fractions containing the blue-coloured product were collected, pooled, lyophilized and stored at −80°C MTfp, NOX4 siRNA (5’-GACCUGACUUUGUGAACAU-3’), scrambled NOX4 siRNA (scramRNA, 5’-GUAAAUUCUCGCGACUAGU-3’) and conjugates thereof were synthesized by Biosynthesis Inc. (Lewisville, USA). MTfp and RNA were produced independently and then chemically conjugated using the crosslinker succinimidyl-4-(N-maleimidomethyl) cyclohexane-1-carboxylate (Supplemental Fig. 4). High performance liquid chromatography and mass spectrometry indicated a purity of greater than 94%. The final products were lyophilized and stored at −80°C.

### 3D Fluorescence Microscopy Imaging

Three sets of 3D fluorescence microscopy experiments were performed. The first set measured the BBB penetration of Cy5 tagged MTfp using DeltaVision Elite integrated with Huygens Deconvolution Software (GE, USA). As an extension of this, a set of sections were prepared to assess the cellular localization of the Cy-5 tagged MTfp in the CNS. The second set of experiments quantitatively assessed the ability of MTfp to direct RNA conjugates to the brain using Leica SP8 X system (Leica, Germany). The third set of experiments addressed the localization of Cy-5 tagged MTfp in various cell types and subcellular objects using the same Leica SP8 X system.

To show that MTfp can cross the BBB, PBS, RVGp-Cy5, revMTfp-Cy5 and MTfp-Cy5 were injected IV into 6-8 week-old, female CD-1 mice (0.5 mM in PBS, 0.1 mL/mouse). Briefly, CD-1 female mice (19.4-25.6 g body weight on day of dosing, 3 per group) were injected once with PBS or one of the peptides. Two hours post injection, mice received a second IV injection of tomato lectin conjugated to FITC (100 µg/mouse, 0.1 mL). Ten minutes later, the animals were terminally anesthetised with an IP injection of 0.2 mL ketamine (200 mg/kg) and Rhompun (20 mg/kg). The bloodstream was flushed by intracardiac perfusion for 10 min at a flow rate of 1 mL/min with heparinized saline (0.9% NaCl, 100 U/mL heparin).

The brains were removed and fixed with 4% paraformaldehyde overnight and then transferred into PBS+0.01% sodium azide and stored at 4 °C. The brains were embedded in 4% agarose, fixed onto the microtome stage and sectioned (20 µm) at 4 °C. The sections were stained with DAPI and then mounted on microscopic slides. Glass coverslips were mounted on the sections using Prolong Gold antifade reagent. In another experiment, to assess the localization of the peptide in the CNS, brains from mice injected with either PBS or Cy5-MTfp followed by FITC-tomato lectin. The brains were collected and processed as stated above. These were then incubated with antibodies targeted against neurons (NeuN; ab177487), microglia (TMEM; ab209064), astrocytes (GFAP; ab7260), lysosomes (LAMP-1; sc-19992-AF546) and endosomes (EEA-1; sc-137130), secondary antibody staining was done for unconjugated primary antibodies with Alexa fluor 680 goat anti-mouse IgG (cat# A21058), Alexa fluor 680 goat anti-rabbit IgG (cat# A21076) and then counter-stained with DAPI, cover slipped, imaged and quantitatively analysed to determine the extent of localization of various cell types and subcellular objects with MTfp. To determine the extent of delivery of RNA to the brain, 6 to 8-week-old female BALB/c mice (16-20 g in weight) received 3 IV injections at 1h intervals with 0.1 mL/mouse PBS, siRNA_AF680_ (30 mg/kg) or MTfp-siRNA_AF680_ (30 mg/kg). Two hours post injection, mice received a second IV injection of tomato lectin conjugated to FITC (100 µg/mouse, 0.1 mL). The mice were then sacrificed and microscope slides were prepared as described above.

In the first set of imaging experiments, 3D fluorescence images were acquired with a DeltaVision Elite integrated with Huygens Deconvolution Software (GE, USA) using a high-resolution Olympus 60X/1.4 or 40X/1.3 Plan-Apochromat oil immersion objective lens. Excitations were performed using solid state illumination Insight SSI. Illumination irradiance controlled by software selectable ND filters (1, 3, 10, 32, 50, 100%T). All images were captured using 12-bit digit CoolSnap HQ CCD camera. The backscattered emission signals from the sample were delivered through the Chroma Sedat Quad ETfilter set. For 3D image data set acquisition, the excitation beam was first focused at the maximum signal intensity focal position within the brain tissue sample and the appropriate exposure times were selected to avoid pixel saturation. The beginning and end of the 3D stack were set based on the signal level degradation. Series of 2D Images for a selected 3D stack volume were then acquired with the appropriate pixel dimensions selected to satisfy the Nyquist sampling criteria. Compiled 3D image data sets were then deconvolved with theoretical point spread functions (or optical transfer functions) using the iterative deconvolution method.

The second and third set of experiments, which quantitatively assessed the ability of MTfp to direct the delivery of a therapeutic compound (NOX4 siRNA) to the brain as well as various localization issues, were performed using Leica SP8 X system (Leica, Germany). 3D confocal images were acquired with a Leica AOBS SP8 laser scanning confocal microscope (Leica, Heidelberg, Germany) using a high-resolution Leica 63X/1.4 or 40X/1.3 Plan-Apochromat oil immersion objective lens. Excitations were performed using either diode or tunable white light laser sources. All images and spectral data (except DAPI) were generated using the highly sensitive HyD detectors (with gated option) in de-scanned mode. The backscattered emission signals from the sample were delivered through the tunable filter (AOBS), the detection pinhole, spectral dispersion prism, and finally to the PMT/HyD detectors. For 3D image data set acquisition, the excitation beam was first focused at the maximum signal intensity focal position within the brain tissue sample and the appropriate HyD gain levels were then selected to obtain the pixel intensities within range of 0-255 (8-bit images) using a color gradient function. The beginning and end of the 3D stack were set based on the signal level degradation. Series of 2D images for a selected 3D stack volume were then acquired with 1024×1024 pixels. The 3D stack images with optical section thickness (z-axis) of approximately 0.3 µm were captured from tissue volumes.

Spectral measurements to confirm the presence of Cy5 signal in the brain parenchyma were also performed using 32-channel Nikon Spectral Detector integrated with Nikon A1 MP+ Multi-Photon Microscope system (Nikon Instruments, New York). The laser used to produce the fluorescence emission from Cy5 was a mode-locked femto-second Spectra-Physics InSight DS femtosecond single-box laser system with automated dispersion compensation tunable between 680-1300 nm (Spectra-Physics, Mountain View, CA). The laser output was attenuated using AOTF and the average power was consistently maintained below the damage threshold of the samples. The power attenuated laser was directed to a Nikon scan head coupled with Nikon upright microscope system (Nikon Instruments, New York). The laser beam was then focused on the specimen through a high numerical aperture, low magnification, long working distance, water immersion objective, CFI75 Apo Water 25X/1.1 LWD 2.0mm WD. The backscattered emission from the sample was collected through the same objective lens. Nikon NIS Element Software was used for the image acquisition.

For each tissue volume reported here, z-section images were compiled and finally the 3D image restoration was performed using IMARIS software (Bitplane, Zurich, Switzerland) or VOLOCITY software (Perkin Elmar, UK). The volume estimation was performed on the 3D image data sets recorded from four or more either cortical or hypothalamus areas of brain tissue samples. Algorithms were developed to automate the quantification of the test articles, blood capillaries and to accurately quantify test articles localized in blood capillaries versus brain parenchyma. In these procedures, a noise removal filter (either Gaussian or kernel size of 3X3) was used to remove the noise associated with the images. To define the boundary between the objects (for instance, blood capillaries) and the background, the lower threshold level in the histogram was set to exclude all possible background voxel values. The sum of all the voxels above this threshold level is determined to be the volume. Fields from each experimental group were pooled. ANOVA was used to assess the volume fractions (for instance, MTfp-Cy5) in the brain parenchyma (Prism 6, Graphpad, La Jolla, USA). The data were log transformed to meet the assumption of homogeneity of variance. Post hoc analysis used a Dunnett’s Test for multiple comparisons. The level of significance was p < 0.05 for the ANOVA and the p-values were corrected for multiple comparisons in the post hoc analysis.

### RNA isolation and real time-PCR

Total RNA was isolated using TRIzol® reagent (Invitrogen, Burlington, Canada) from frozen brains of mice that had been with PBS, NOX4 siRNA, MTfp-NOX4 siRNA or MTfp-scramRNA. RNA was reverse transcribed to cDNA with a Script cDNA SuperMix agent(QuantaBio, Gaithersburg, USA) using random hexamers and oligo dT. SYBR^®^ Green PCR Master Mix and RT-PCR Reagents kit (Applied Biosystems, Foster city, USA) were used to do real time PCR on an ABI 7500 FAST REAL TIME PCR System (Applied Biosystems, Foster city, USA). Nox4 primers (5’-CTTGACTTGATGGAGGCAGTAG-3’and 5’-GCCTTTATTGTGCGGAGAGA-3’) and GAPDH primers (5’-AACTTTGGCATTGTGGAAGG-3’ and 5’-ACACATTGGGGGTAGGAACA-3’) were used.

The master reaction mixture consisted of 1X SYBR®Green PRC buffer, 3 mM MgCl_2_, 1 mM dNTP, 0.625 U Taq polymerase, 0.25 U Amperase UNG, 10 ng cDNA and 300 nM primers (sense and antisense). The real time PCR was carried out at 50 °C for 2 min, 95 °C for 10 min, followed by 40 cycles at 95 °C for 15s, and 60 °C for 1 min. The real time PCR efficiency was determined for each gene with the slope of a linear regression model, and all samples displayed efficiencies between 84% and 96%. Negative controls containing no target RNA were included with each of the real time RT-PCR runs. Data are expressed as mean ± standard error of mean. In multiple group comparisons, data were analyzed by one-way ANOVA with Tukey post-hoc test. *P*-values less than 0.05 were considered significant.

### Stroke induction by transient middle cerebral artery occlusion

Work with the mouse stroke model was performed at the NRC (Ottawa, Canada) and the Centre for Comparative Medicine (University of British Columbia, Vancouver, Canada). Male C57Bl/6 mice (10 weeks old) were injected IV three times, at 1-hour intervals, with PBS, NOX4 siRNA, MTfp-siRNA or MTfp-scram RNA (30 mg/kg). One hour after the last injection, an ischemic stroke was induced on the left hemisphere by MCAO. The external and internal carotid arteries (ECA and ICA, respectively) were dissected and isolated and the common carotid artery (CCA) ligated. A nylon thread coated with dental silicon (Cat. # 60231012, Doccol Corp) was inserted into a small hole in the ECA towards the bifurcation of the CCA. The thread was gently introduced into the ICA, passed into the Circle of Willis thus occluding the opening of the middle cerebral artery. Body temperature was not allowed to drop below 36°C and anesthesia was maintained throughout the surgery occlusion. The filament was removed after 1h, and the ligature on the CCA was removed allowing reperfusion of the affected brain area. Mice were sacrificed at 24 hrs post reperfusion. Sham surgeries (arteries isolated but not occluded) on animals that received all 3 injections of PBS were also performed as controls.

### Stroke Analysis

An individual, who was blinded to the treatment each mouse had received, assessed post stroke behaviour. Mice were assessed at 0.5 hr and 24 hr after recovery from anesthesia using a method described by Jiang *et al*^39^. A score of 0 is normal; 1 is mild turning behaviour with or without inconsistent curling when picked up by tail and <50% attempts to curl to the contralateral side; 2 is mild consistent curling with >50% attempts to curl to contralateral side; 3 is strong and immediate consistent curling, mouse holds curled position for more than 1–2 s, the nose of the mouse almost reaches tail; 4 is severe curling progressing into barrelling, loss of walking or righting reflex; 5 is comatose or moribund. According to this method, the neuroscore at 0.5 hr is a measure of surgical success and identifies animals that had received abnormally severe or mild damage during the procedure. Indeed, mouse #6, belonging to the PBS group, showed a neuroscore of 0, whereas all the other mice in PBS and siRNA pretreated groups exhibited neurorscores between 3.0-3.5. This indicates that the MCAO of mouse #6 was insufficient to induce stroke. The opposite was true of mouse #14 from the MTfp-siRNA treated group, which had a neuroscore of 4.5 indicating that MCAO had caused severe surgical trauma. All the other mice in that group had neurorscores between 1 and 2.5. Data from these two aberrant animals were excluded from final calculations.

To determine infarct size, mice were sacrificed after the second behavioural assessment (24h post-op). Mice were terminally sedated with 0.2 mL of ketamine (200 mg/kg) and Rhompun (20 mg/kg) and perfused transcardially at 1 mL/min for 10 min heparinized saline (0.9% NaCl, 100 U/mL heparin). Brains were removed, sectioned and stained with 2% TTC in PBS to visualize the infarcts. Images of the brain slides were obtained after 15-20 min staining at room temperature. The brain slices were cut in half to separate the ischemic hemisphere and the contralateral hemisphere. Tissue was rinsed with physiological saline and then exposed to a mixture of ethanol/dimethyl sulfoxide (1:1) for 24 hrs in the dark to solubilise the coloured formazan product. The absorbance at 485 nm was measured. Percentage loss in brain TTC staining in the ischemic side of the brain was compared with the contralateral side of the brain of the same animal using the following equation.

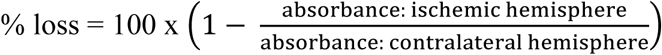

Data are expressed as mean ± standard error of mean. In multiple group comparisons, data were analyzed by one-way ANOVA with Tukey post-hoc test. *P-*values less than 0.05 were considered significant.

## Supplemental Information

Supplemental Information is linked to the online version of this paper at nature.com/natbiomedeng

## Acknowledgements

We thank Professor Lawrence McIntosh from University of British Columbia, Department of Chemistry for access to the NMR equipment, consultation on the NMR experiments and data analysis; Professor Terry Pearson from the University of Victoria, Department of Biochemistry for his comments on the manuscript and for providing the isotopically labelled peptides; Mr. Wade Edris for his technical assistance with confocal and deconvolution microscopy imaging. We also thank MRM Proteomics, Cellial Technologies, the National Research Council of Canada and Ava McHugh for their professional services.

This research is made possible by donations to the Jefferies Laboratory through the Sullivan Foundation at Vancouver General Hospital. Funding for this work was provided to WAJ in the Michael Smith Laboratories, at the University of British Columbia and the Vancouver Prostate Centre, at Vancouver General Hospital, by grants from the Canadian Institutes for Health Research (MOP-133635), the Natural Sciences and Engineering Research Council of Canada (CRDPJ 452456-13) and a University of British Columbia, industrial partnership agreement with Bioasis Technologies Inc. (BTI). TA was supported by NIH grants 1S10OD010756-01A1 and 1S10OD018124-01A1. BAE was supported by a Postdoctoral Fellowships from the Michael Smith Foundation for Health Research, the Centre for Blood Research at the University of British Columbia and the Pacific Alzheimer’s Foundation. CSBS was supported by an Alzheimer’s Drug Discovery Foundation Outstanding Young Investigator Scholarship, and by a William and Dorothy Gilbert Graduate Scholarship in Biomedical Sciences at the University of British Columbia and a scholarship from the Centre for Blood Research Graduate Student Award at the University of British Columbia. The funding sources had no role in the study design, data collection, analysis or interpretation of data, or in the writing of the paper.

## Author Contributions

All authors contributed to the paper and BAE, CSBS and WAJ wrote the manuscript and all authors revised it and approved the final version. WAJ conceived of the project.

## Author Information

Reprints and permissions information is available at nature.com/reprints. The authors declare competing financial interests: MMT, TZV and RG are, or were, employees of BTI. WAJ was a consultant and the founding scientist of BTI. BTI contact regarding xB^3 TM^: Dr. Mei Mei Tian, Vice President, Head of External Research, Bioasis Technologies Inc., 14 Water Street, Guilford, CT 06437, USA

## Supplemental Information

**Supplementary Table 1.**
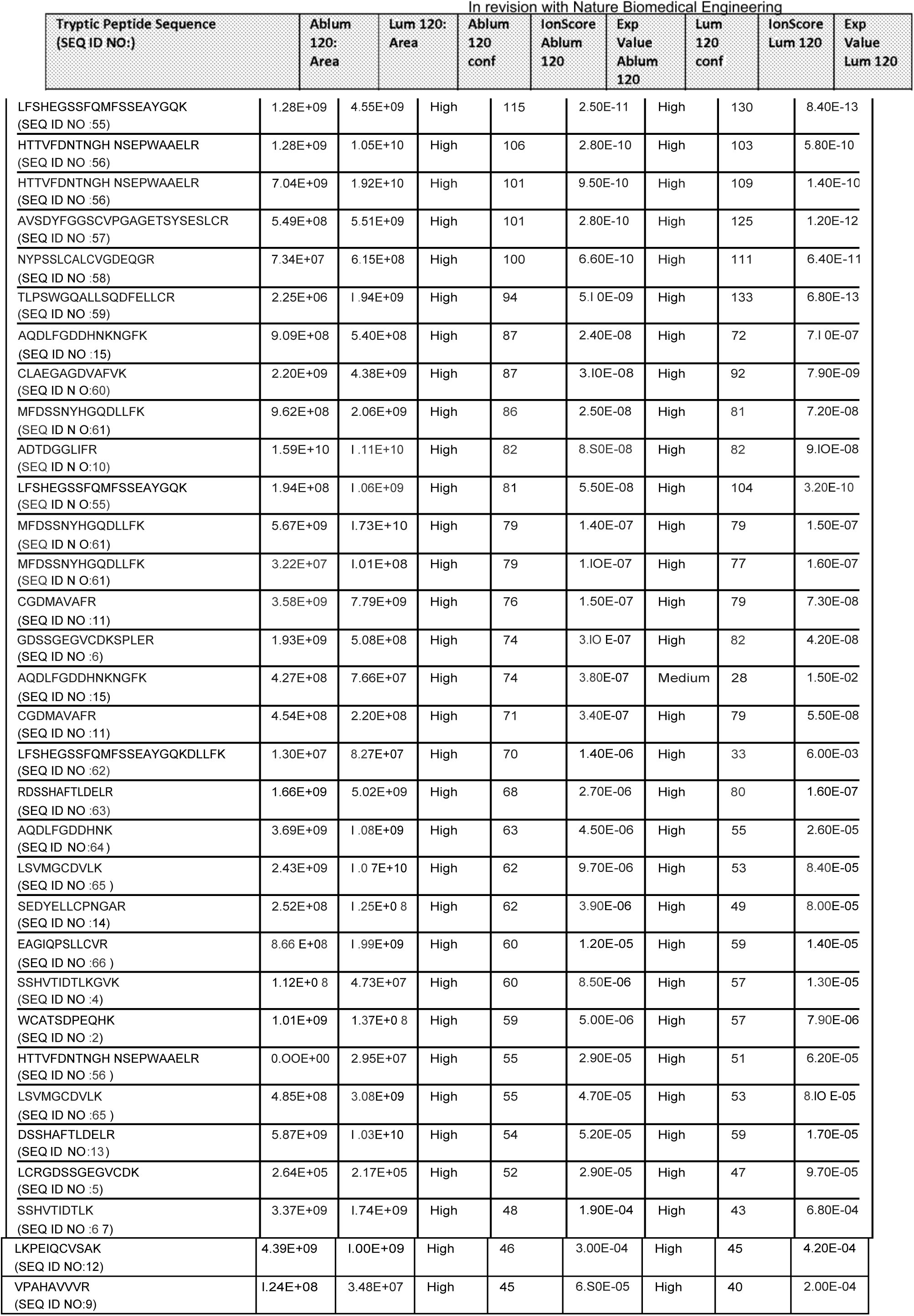
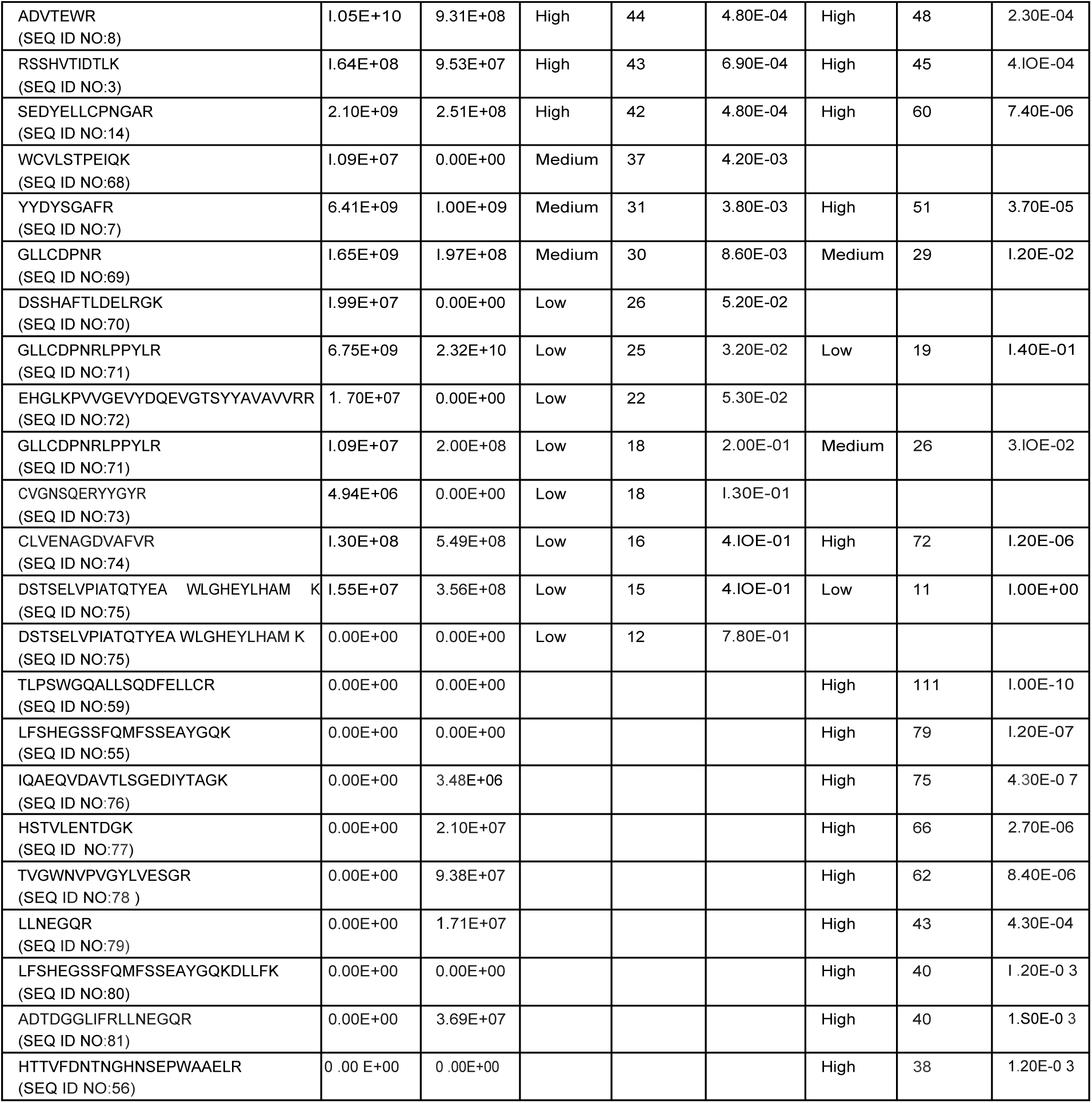
Tryptic Peptides at 3 Micron Pore Size

**Supplemental Table 2.**
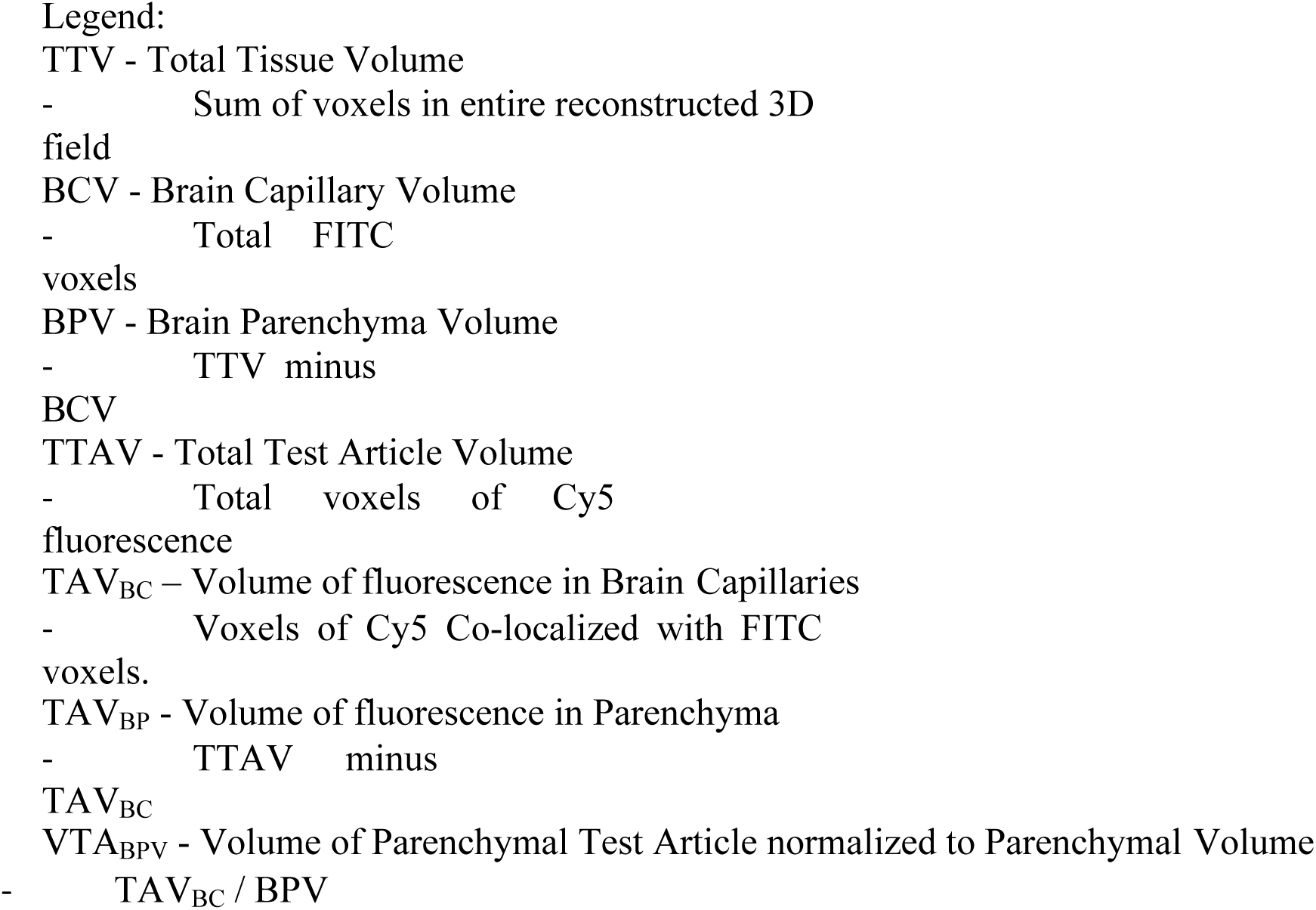
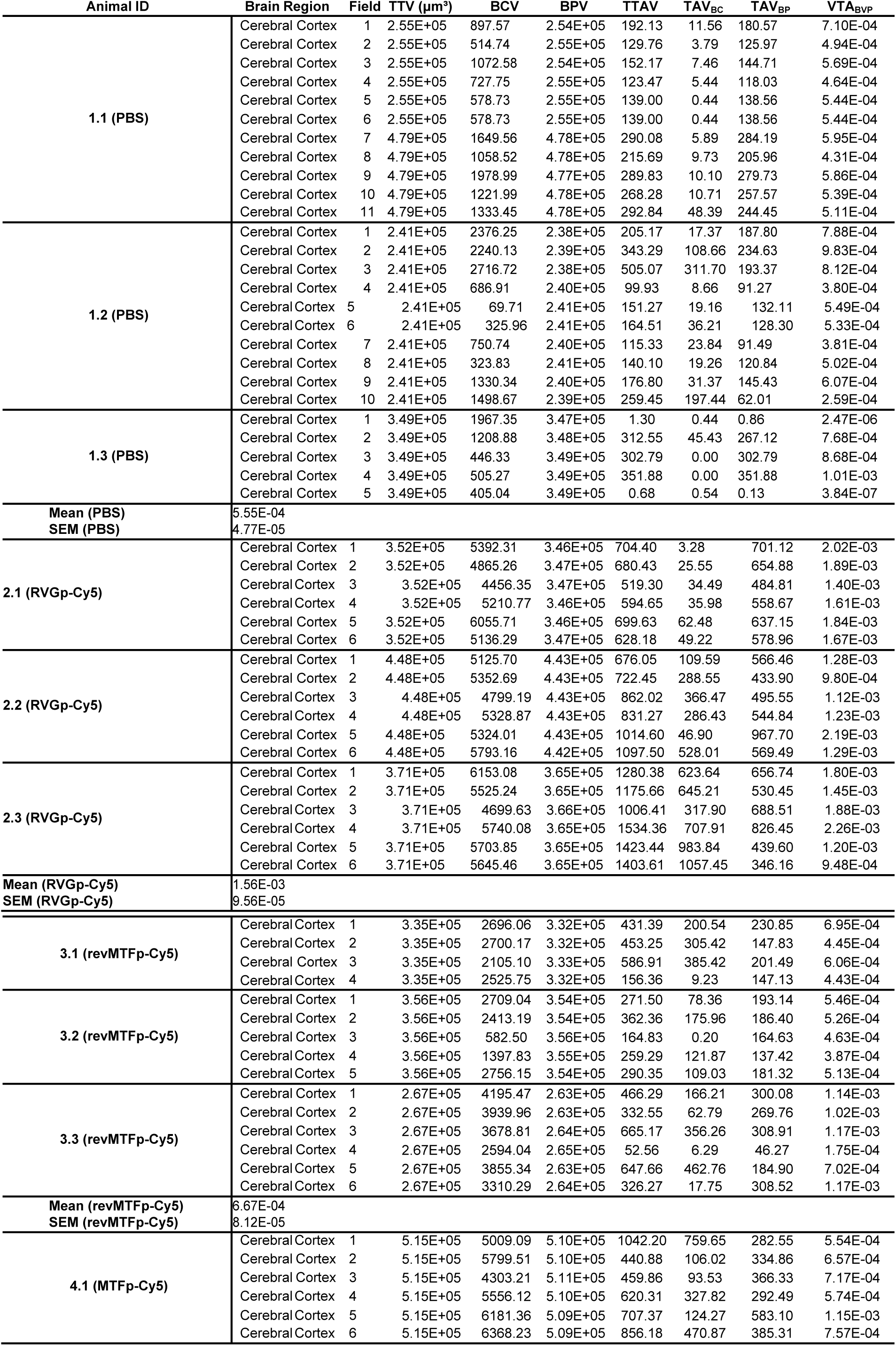
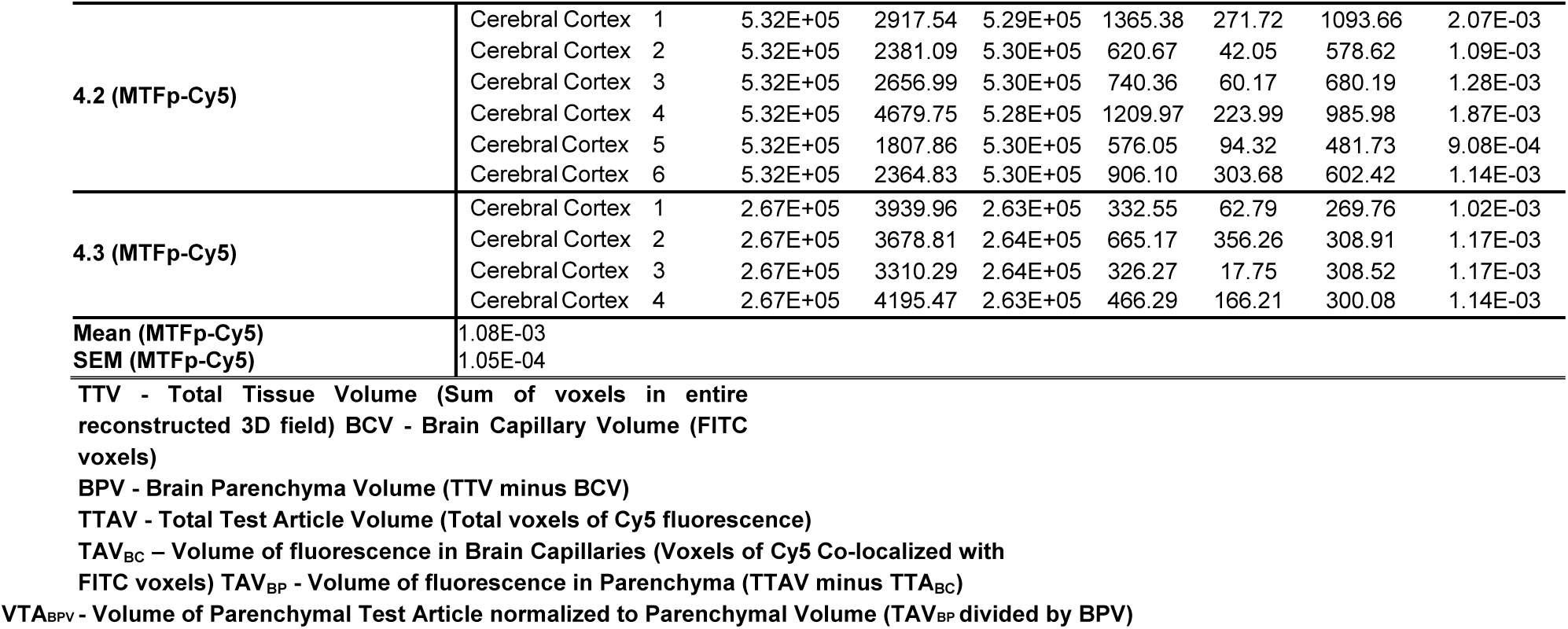
Fluorescence associated with localization of MTfp to brain tissues.

**Supplemental Table 3.**
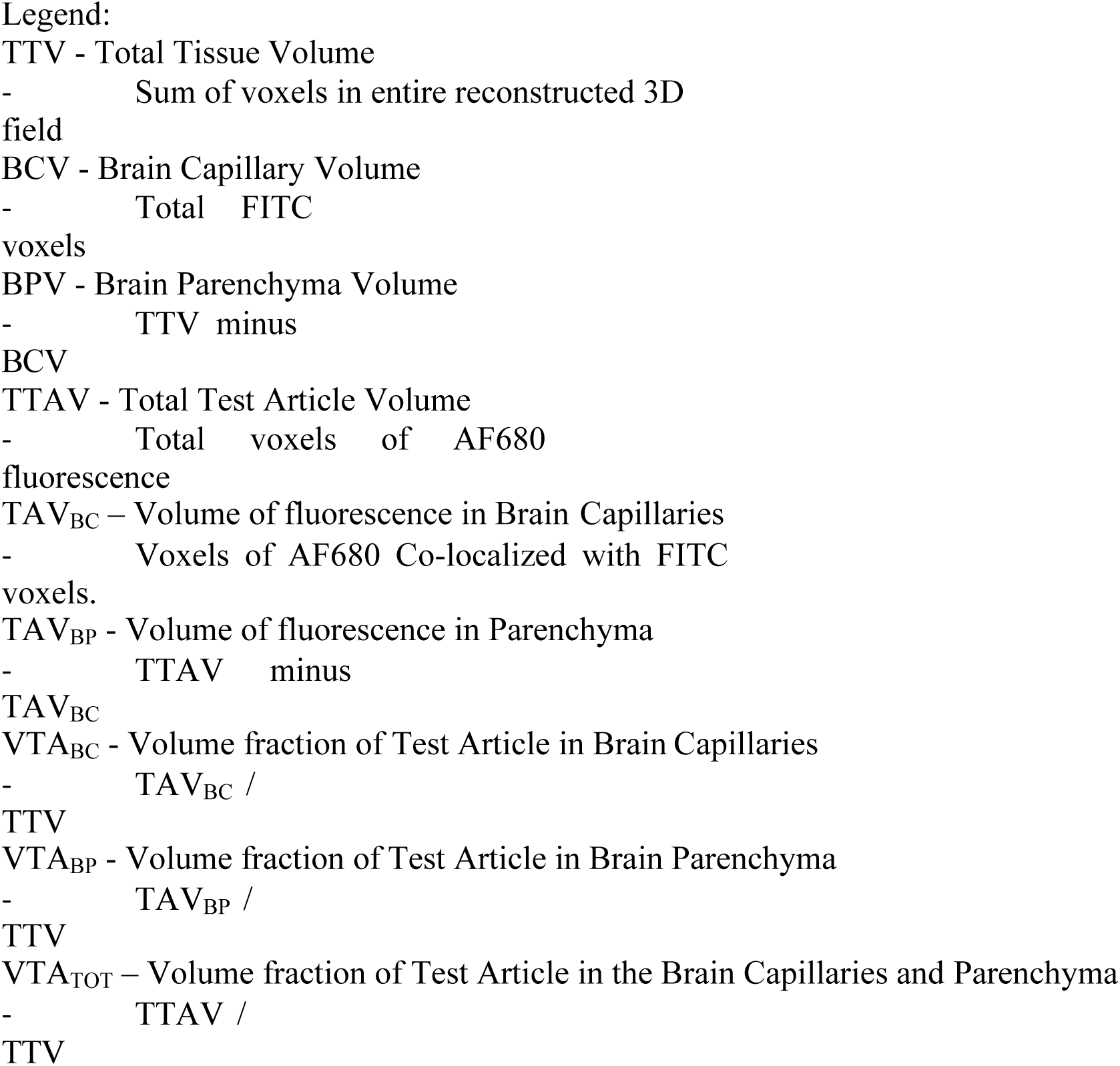
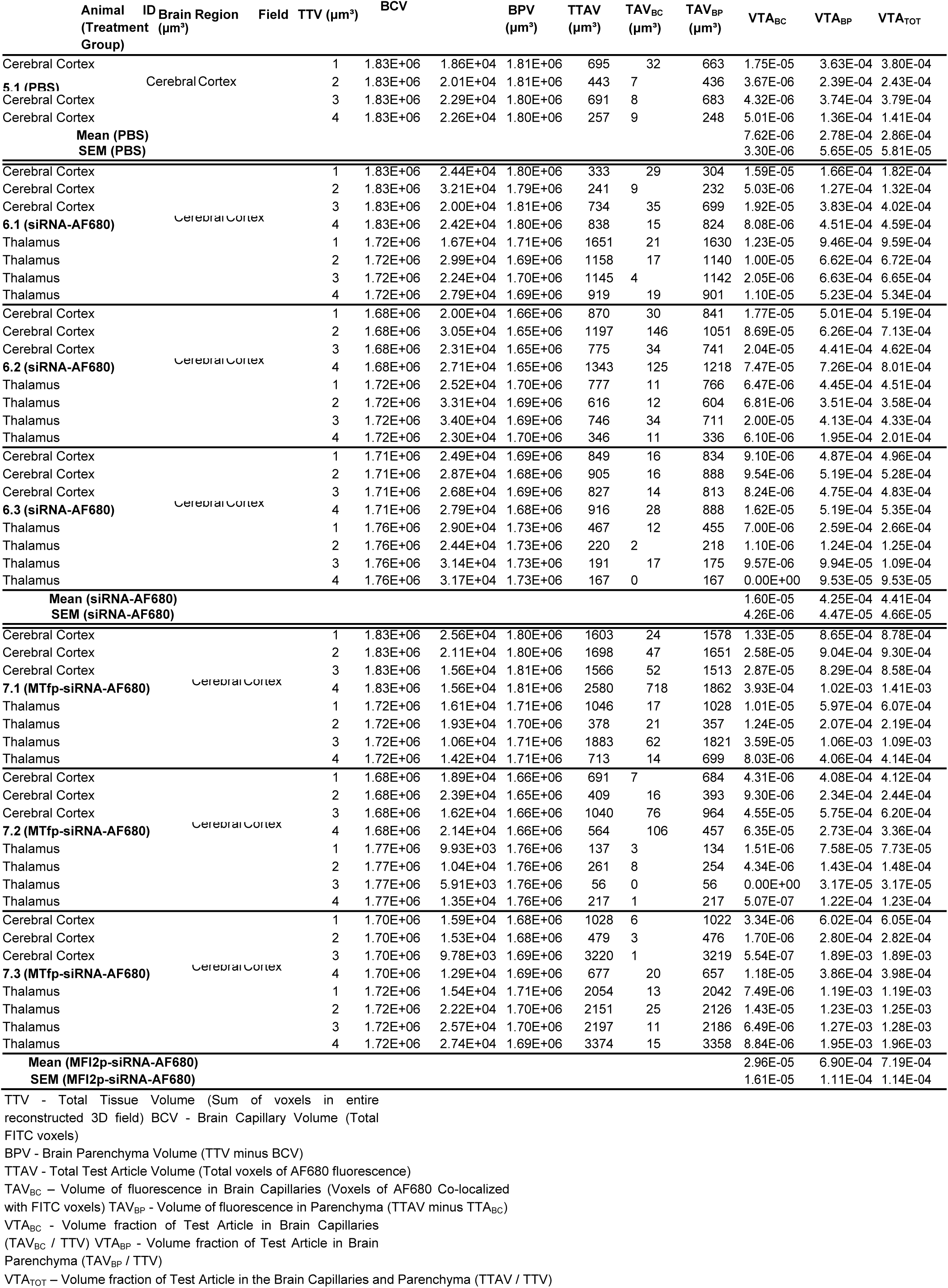
Fluorescence associated with the localization of siRNA and MTfp-siRNA to brain tissues.

**Supplemental Table 4.**
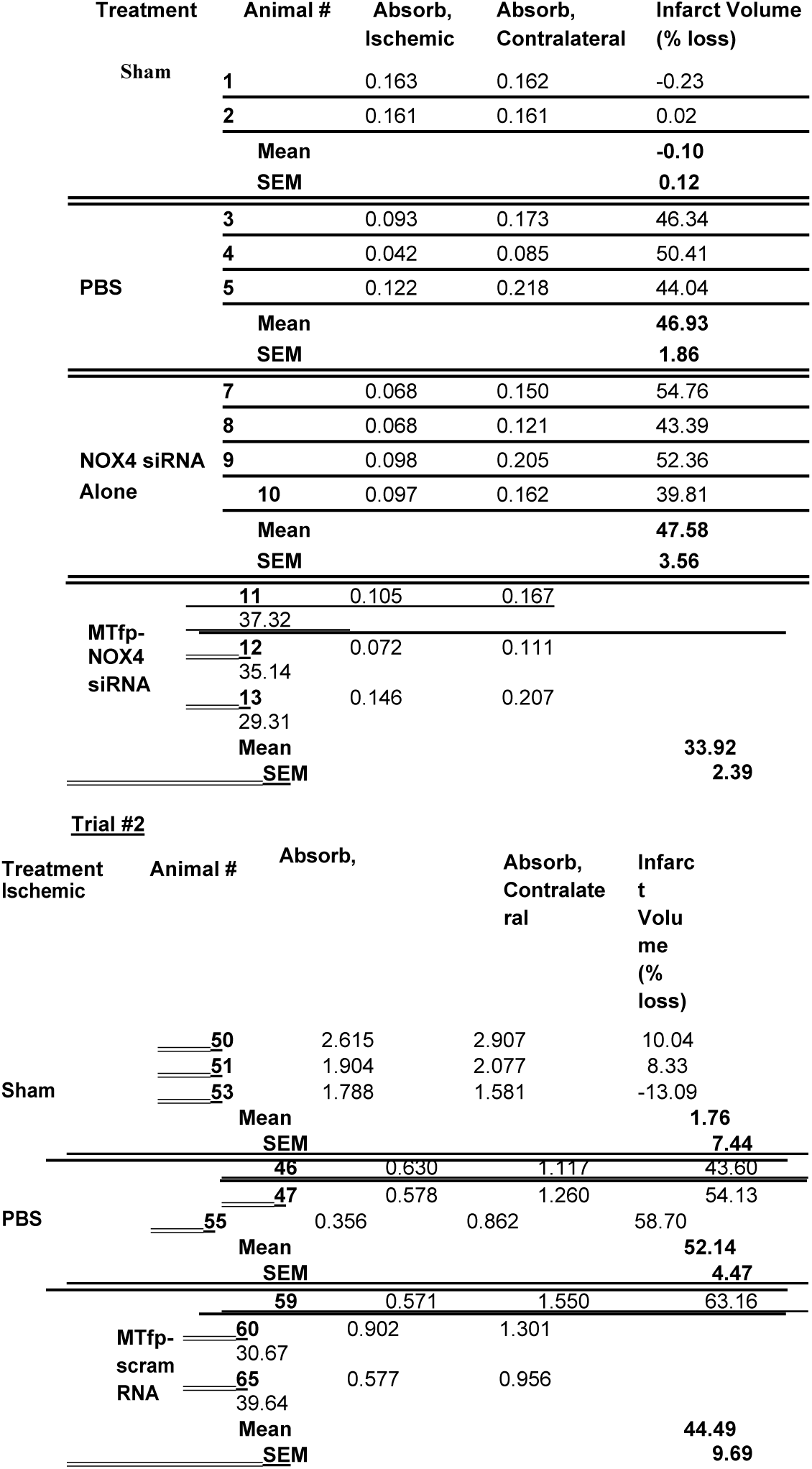
Quantitation of stroke infarct size by TTC assay Trial #1

**Supplemental Figure 1.**
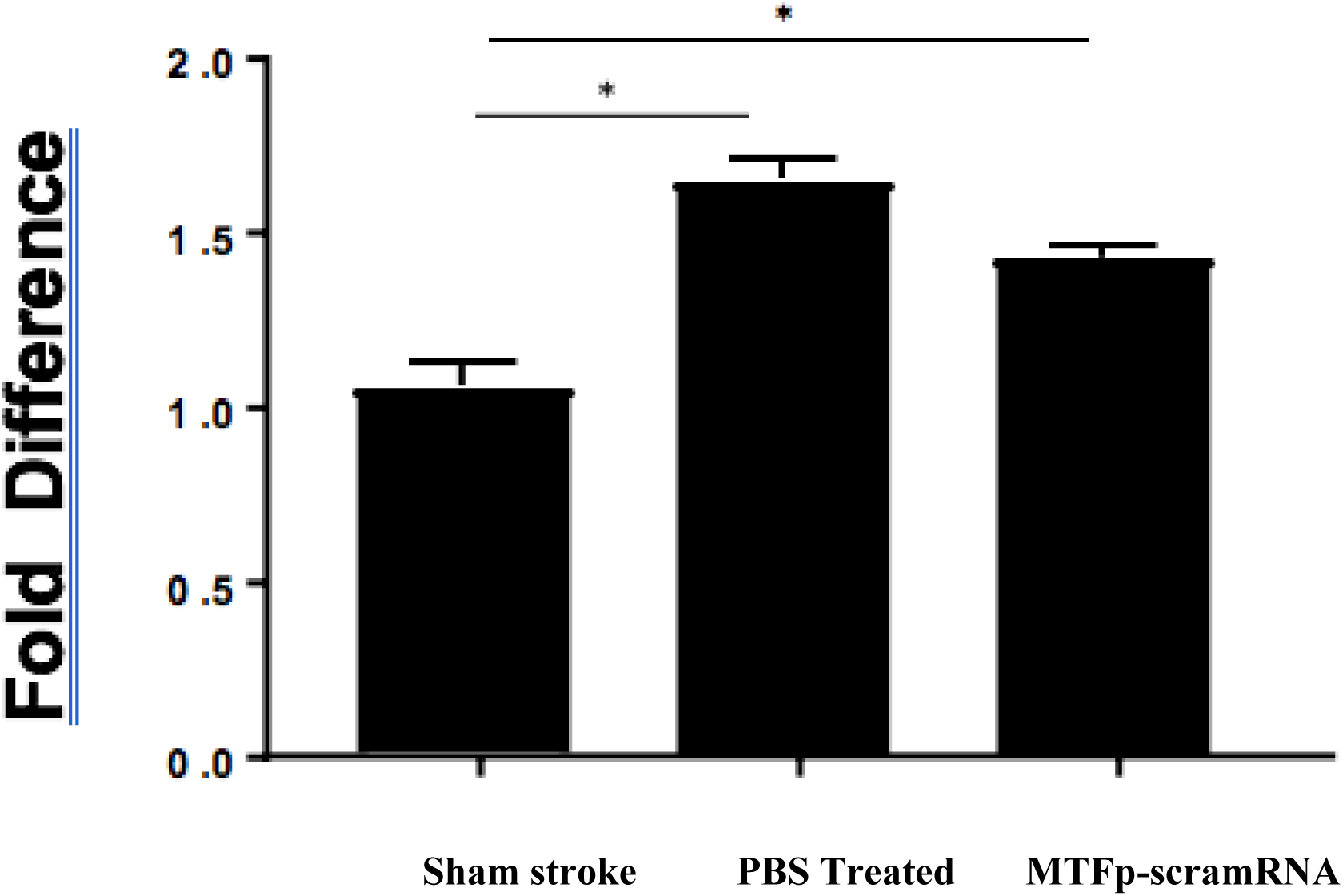
NOX4 mRNA expression in the brain after MTfp-scramRNA treatment and stroke induction. Relative gene expression of *Nox4* in the mouse brain hemispheres by RT-PCR following treatment and 24 hours post-stroke (GAPDH as a reference gene). For each treatment group, the *Nox4* expression in the left hemisphere (stroke) is normalized against the right hemisphere (contralateral) of the same animal. Data are shown as mean ± SEM.

**Supplemental Figure 2.**
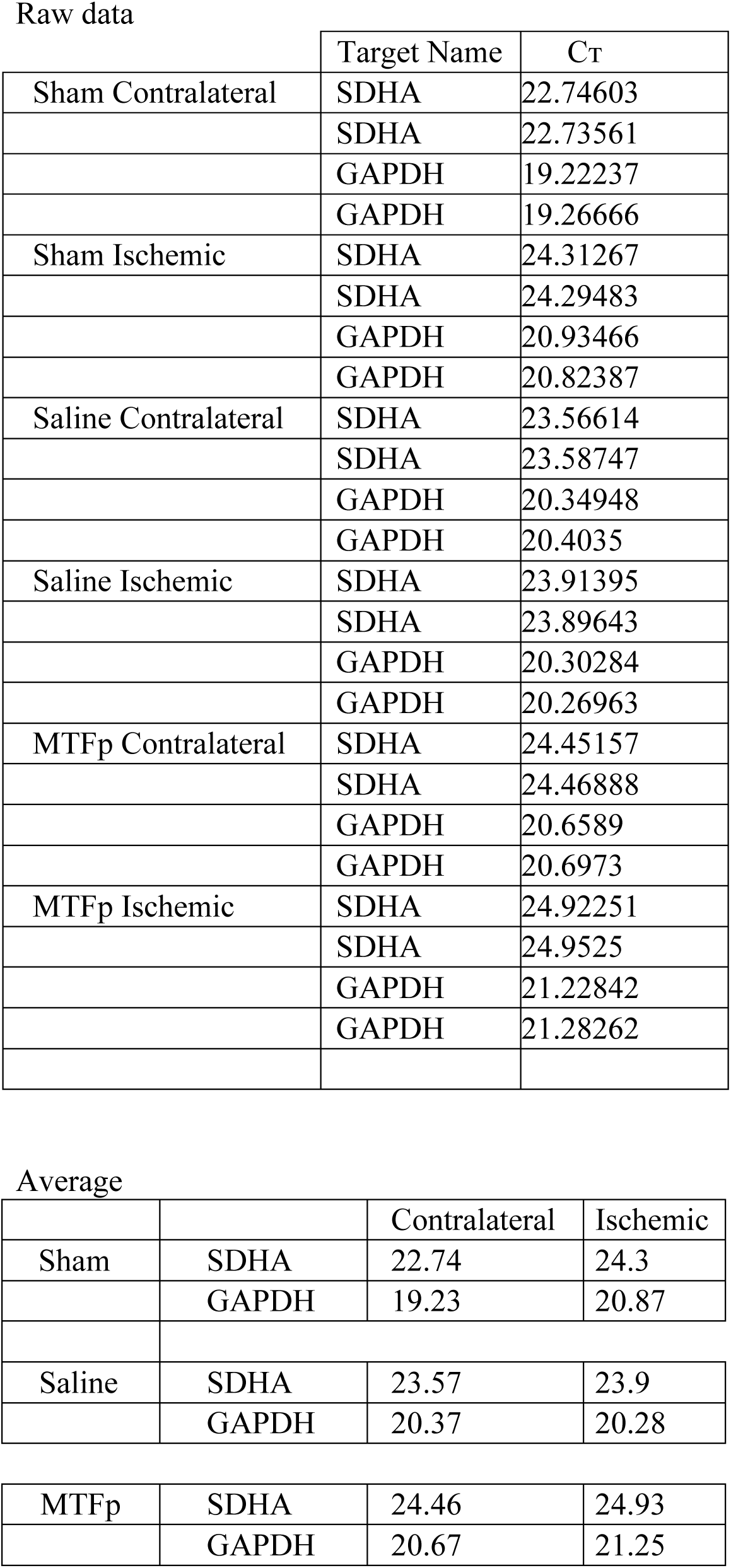
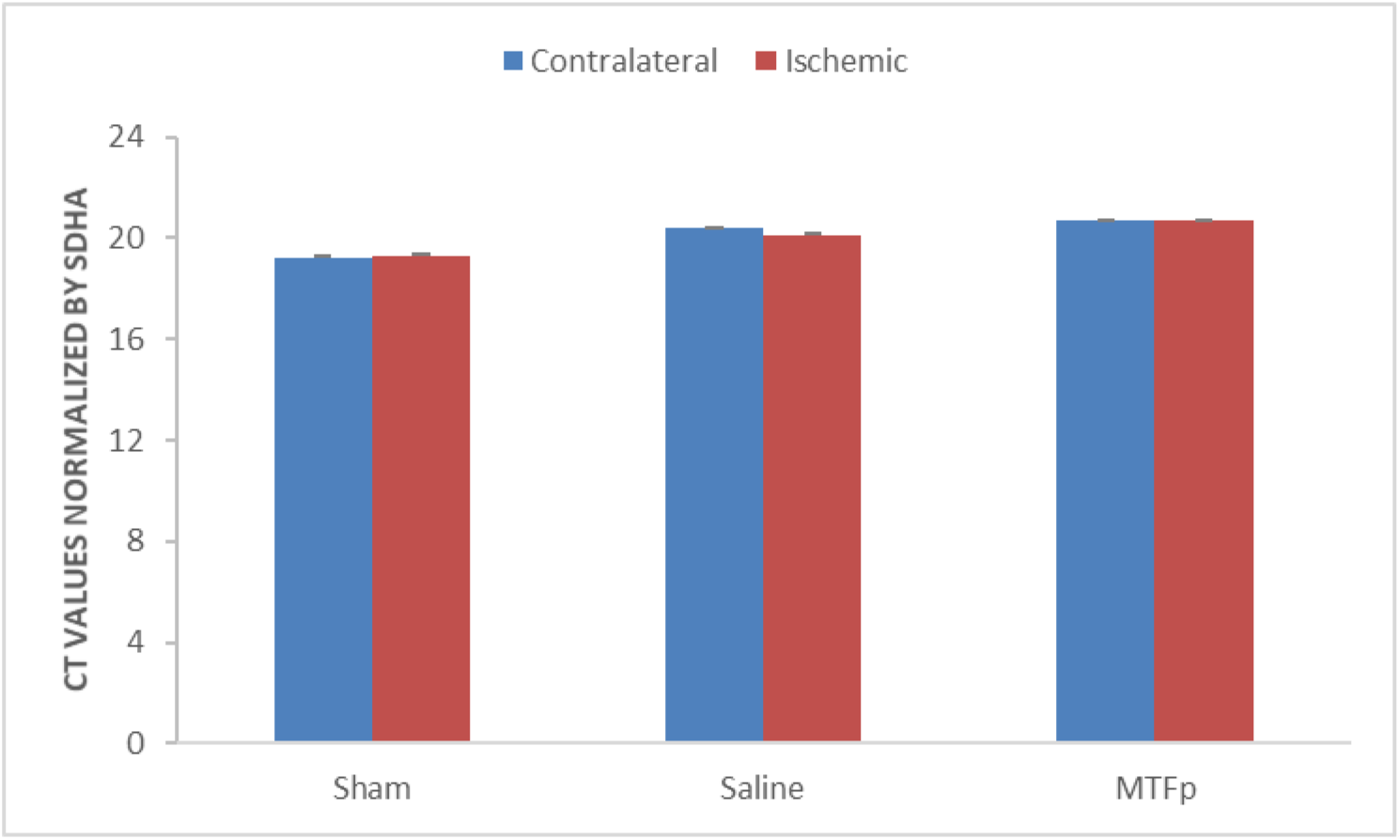
GAPDH as a reliable reference gene to evaluate NOX4 mRNA expression. GAPDH expression was normalized to succinate dehydrogenase complex flavoprotein subunit A (SDHA) expression, a gene known to be a reliable reference gene in brain ischemia. Sham = surgical control (no hypoxia). Saline = surgical hypoxia induction with saline injection. MTFp = surgical hypoxia induction with MTF peptide injection. Error bars represent standard error of mean of duplicate values. GAPDH gene expression does not significantly change after induction of ischemia in the brain, making it a reliable reference gene to evaluate NOX4 mRNA expression.

### Method

Frozen brain tissue samples were first processed with 70 7m cell strainer to disintegrate them and immediately transferred to lysis buffer (Qiagen) and then homogenised using an 18G needles and syringes. RNA was isolated using the RNeasy^®^ plus Mini Kit (Qiagen) in accordance with the manufacturer’s instructions Subsequently RNA was quantified spectrophotometrically (NanoDrop, ThermoFisher Scientific) at 260 nm (A_260_) and purity was estimated by an A_260_/A_280_ ratio > 1.8. The integrity, purity, and amount of RNA were verified by visualization of total RNAs after electrophoresis on 1% agarose gel. 2 µg of RNAs were reverse transcribed using a SuperScript II Reverse Transciptase (Thermofisher Scientific) according to the manufacturer’s instructions.

RT-Q-PCR reactions were carried out for GAPDH and SDHA in each sample using GeneAmp 9600 Fast Instrument (ThermoFisher Scientific). Each reaction was performed in a final volume of 20 µL containing 2µL cDNA diluted with H_2_O, 10 7l of iTaq Universal SYBR® Green Supermix (Bio-Rad) and 17l of each gene specific primers (SDHA Fwd primer: 5’-GGAACACTCCAAAACAGACCT 3 and Rev primer” 5’-ACCACTGGGTATTGAGTAGAA-3’, GAPDH Fwd primer: 5’-AACTTTGGCATTGTGGAAGG-3’ and Rev primer 5’-ACACATTGGGGGTAGGAACA-3’. All samples were run in triplicate and average values were calculated. Two independent reverse transcriptions were tested for each gene.

**Supplemental Figure 3.**
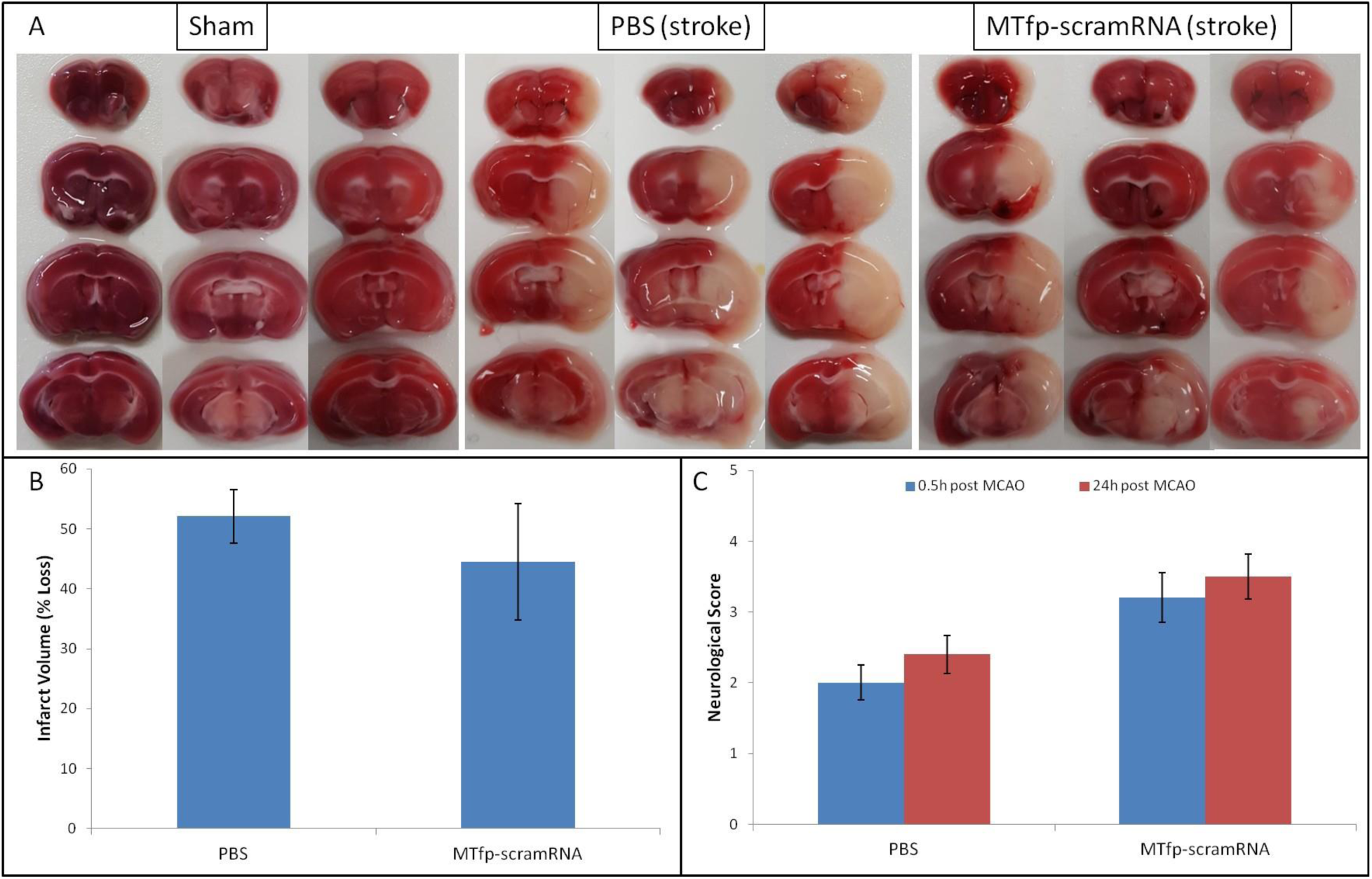
Treatment with MTfp-scrambled RNA does not offer neuroprotection after ischemic stroke. [A] TTC-stained brain sections of mice receiving various IV injections, followed by ischemic stroke. [B] Infarct volume was quantitated by measuring absorbance (485nm) of solvent extracted dye. Data are shown as mean ± SEM (no significant difference). [C] Neuroscore at 0.5 hr and 24 hr after stroke induction. Data are shown as mean ± SEM (There is no significant difference between the treatments at either time point).

**Supplemental Figure 4.**
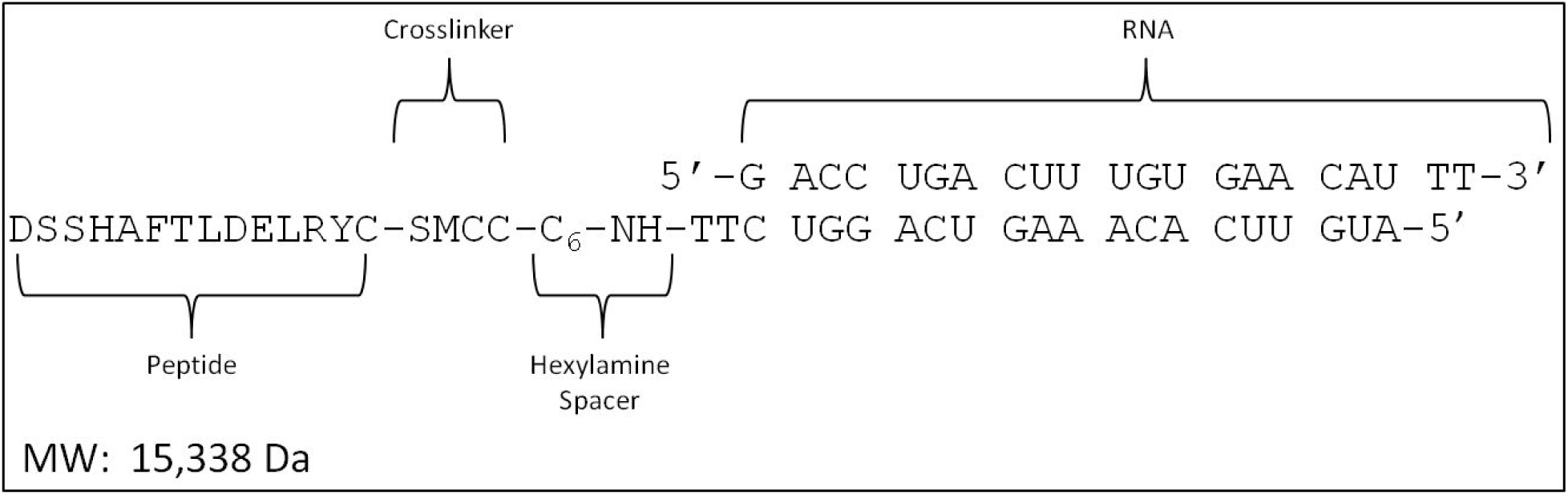
Design of the MTfp-RNA conjugates. SMCC = succinimidyl 4-(N-maleimidomethyl) cyclohexane-1-carboxylate.

## Supplemental Discussion 1. Structural Characterization of MTfp and elaboration of a candidate receptor

### Structural Characterization of MTfp

Structural studies were carried out to identify unique features of MTfp that enable it to retain the ability of the parent molecule to cross the BBB and also differentiate it from its phylogenetic cousins such as Tf and Lf, which do not efficiently cross the BBB. MTfp (DSSHAFTLDELR) is well conserved through evolution (Supplemental Discussion Table 5). There is perfect sequence identity among most primates and strong conservation among many other mammals. Divergence is typically restricted to the first five residues (DSSHA). When there is a mutation, the altered amino acid is often physiochemically similar and therefore, potentially benign. The lack of variability in the C-terminal portion of MTfp suggests that this region is most important for receptor binding. This supposition is supported by the facts that human MTfp was identified by transcytosis across a model of the bovine BBB and *in vivo* transcytosis is observed in mice. Both bovine and murine MTfp homologues differ from the human version by two amino acids; both near the N terminus (Supplemental Discussion Table 1). MTfp shows very little primary sequence identity with analogous peptides from the close MTf relatives, Tf or Lf (Supplemental Table 4), this feature potentially explaining MTf’s unique ability for BBB transcytosis compared to other members of the transferrin family.

The crystal structure of MTf is unknown but *in silico* homology models were generated using the Robetta structure server^1^ (data not shown) and compared to published crystals structures of human Tf^2^. Although the primary amino acid sequence of MTfp and the analogous Tf peptide (Tfp) are different, their location on their respective proteins and their secondary structures are predicted to be nearly identical. Both are displayed as surface-exposed amphipathic turns/loops between beta-strands. The hydrophobic residues of MTfp (Phe_465_, Leu_467_ and Leu_470_) are oriented towards the core of the protein, analogous to hydrophobic residues in Tfp, while the remaining residues appear to be solvent-exposed. In its protein context, both MTfp and Tfp are generally unstructured but anchoring by the hydrophobic residues seems to impart a slight helical character in the conserved C-terminal region (FTLDELR). Based on secondary structure comparison, it appears that the key differences between MTfp and Tfp are MTf His_463,_ which is absent in the shorter, 11 AA Tfp and the substitution of an asparagine in Tfp for glutamate in MTfp at position 469 in the putative helical region. It is likely that the polar orientation and slight helical conformation of MTfp are important for MTfp’s receptor binding capability. In the MTf protein context, this polarity would be maintained by repulsion between adjacent acidic residues and hydrophobic interactions with the core of the protein. However, these stabilizing hydrophobic interactions are absent when MTfp exists as a peptide in solution. It is possible that an entropic cost, needed to orient the free peptide into an appropriate receptor-binding conformation, may reduce receptor affinity and thus rates of transcytosis. However, we have shown that MTfp is capable of transcytosis, comparable to intact MTf. Since this is the first known example of a peptide, derived from a larger plasma protein, being able to cross the BBB, we chose to assess the molecular dynamics of this peptide in solution over a range of temperatures (10°C, 20 °C, 30 °C and 37 °C) by proton-carbon heteronuclear multiple bond correlation spectroscopy (HMBC) and proton-proton Nuclear Overhauser Effect Spectroscopy (NOESY) nuclear magnetic resonance (NMR). As expected, based on chemical shift data from the HMBC experiments, at all temperatures, MTfp is calculated to be to be an unstructured coil (data not shown). However, at cooler temperatures (10 °C and 20 °C), amide to amide NOESY signals, consistent with a helical conformation, were also observed (Supplemental Discussion Supplemental Figure 5). Secondary structure is unusual in such a small peptide; however, our experimenta data suggests that MTfp has the propensity to adopt a transient helical character, possibly driven by repulsion between the neighbouring acidic residues. Perhaps this tendency towards a helix, similar to its conformation while part of MTf, reduces the energetic cost of peptide-receptor binding and contributes to the peptide’s unique transcytosis potential.

### MTf Receptor Candidate

A broadly-specific receptor (low-density lipoprotein receptor-related protein) is capable of binding plasminogen-MTf complexes^3^;however this receptor does not appear to bind free, soluble MTf. We speculate that the protein, inactive N-acetylated alpha linked acidic dipeptidase like 2 (NAALADL2; UniProtKB - Q58DX5) may be a candidate MTf/MTfp receptor on the BBB. NAALADL2 is a type I transmembrane protein and is distantly related to the Tf receptors but shares more sequence conservation with the Tf receptors than it does with the other members of the NAALAD family. Members of this family are known to bind and cleave variations of an aspartate-glutamate motif. This dipeptide sequence is found within MTfp (DSSHAFTLDELR). Owing to a collection of point mutations, NAALADL2 is predicted to be, proteolytically inactive. Furthermore, similar to the Tf receptors, and unlike the other NAALADs, NAALADL2 possessed a tyrosine-based endocytosis signaling motif in its cytoplasmic domain; a necessary feature if it were to mediate transcytosis. Finally, NAALADL2 is distributed in tissues where MTf is known to accumulate (ProteinAtlas). We hypothesize that NAALADL2 is a specific MTf receptor.

**Supplemental Discussion Table 5.**
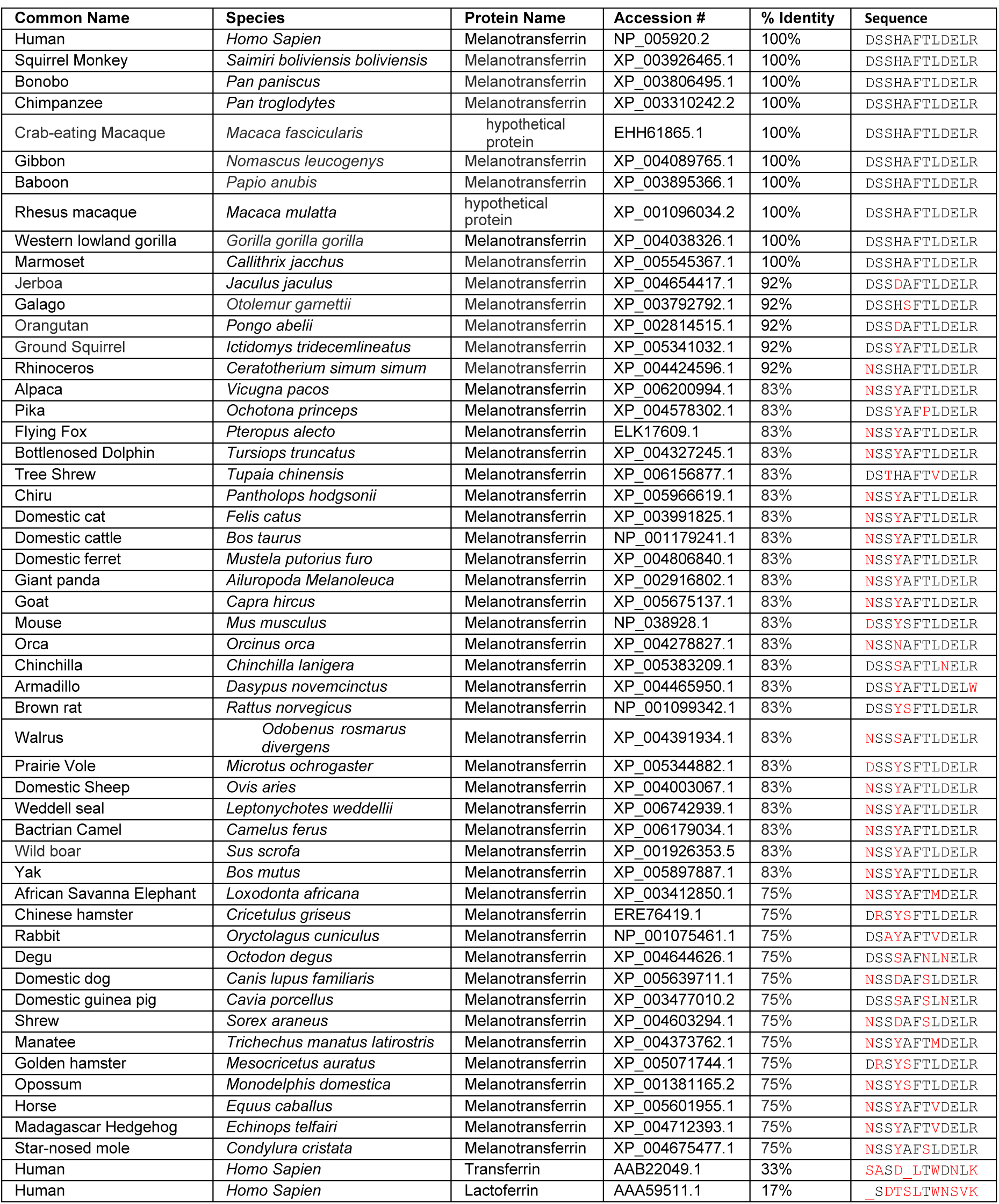
Phylogenetic comparisons of MTfp homologues

**Supplemental Figure 5.**
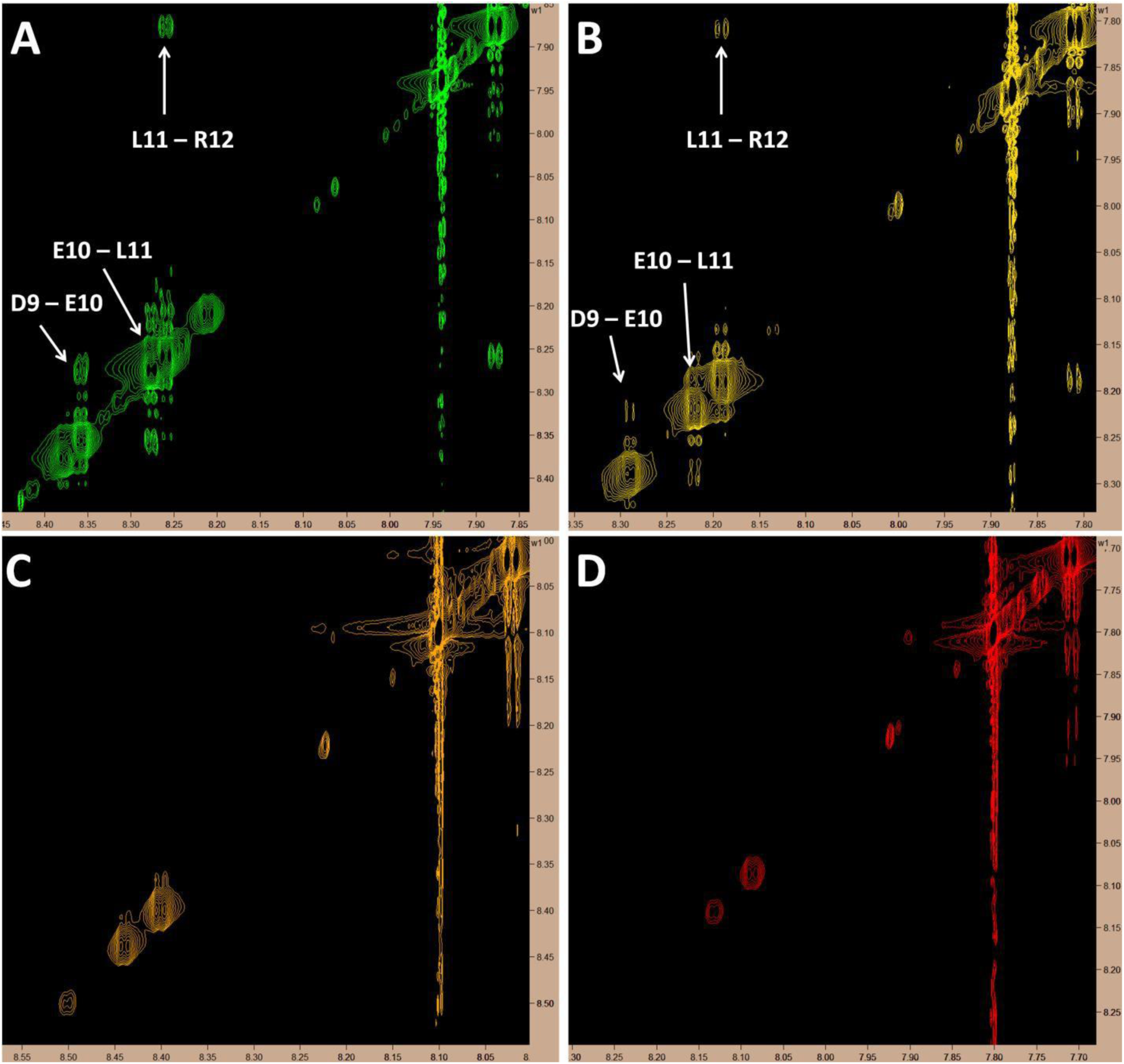
Proton NMR showing amide to amide proximity. Cross-peak signals suggestive of a helical structure in the C-terminus of MTfp can be seen at 10 °C and 20 °C but all signals in this region fade as temperature rises; possibly due to proton exchange with solvent. All images represent 5 mM MTfp in 25 mM sodium phosphate buffer at pH 7 with 5% D_2_O. **(A)**: 10 °C. **(B)**: 20 °C. **(C)**: 30 °C. **(D)**: 37 °C.

## Methods

### Nuclear Magnetic Resonance

All experiments were performed in a Bruker Ascend 850 MHz magnet. Data were processed and analyzed using Bruker’s TopSpin and Sparky^5^ respectively. Proton correlation (COSY), total correlation (TOCSY), nuclear Overhauser effect (NOESY) spectroscopy as well as proton-carbon heteronuclear single quantum correlation (HSQC) and heteronuclear multiple-bond correlation (HMBC) spectroscopy were performed on synthetic, 5 mM DSSHAFTLDELR peptide in 25 mM sodium phosphate buffer (pH 6.5) with 5% D_2_O at 10 °C, 20 °C, 30 °C and 37 °C. Spectral assignment was performed manually and chemical shift data were assessed by MICS (Motif Identification from Chemical Shifts) software.

